# Reg3β removes aged neutrophils after myocardial infarction

**DOI:** 10.64898/2026.02.19.706868

**Authors:** Holger Lörchner, Laia Cañes Esteve, Julia Detzer, Roxanne Harzenetter, Lucia Camacho Pulido, Maria Elisa Góes, Christian Wächter, Sophia Schlattner, Stefan Günther, Carsten Kuenne, Mario Looso, Yousef Alayoubi, Filip Klicek, Maja Pučić-Baković, Gordan Lauc, Diana Campos, Miloslav Sanda, Matthias Gunzer, Matthias Totzeck, Jochen Pöling, Thomas Braun

## Abstract

Recruitment and resolution of immune cell accumulation after myocardial infarction (MI) is critical for effective wound healing and prevention of maladaptive remodeling. Numerous signals are known to recruit neutrophils, critical drivers of inflammation, but knowledge about signals containing and resolving their accumulation is limited. We discovered that Regenerating-islet derived protein 3 beta (REG3β) limits the persistence of neutrophils after MI, thereby promoting resolution of inflammation. REG3β selectively binds to aged and hyperactive neutrophils and induces rapid cell death, enabling clearance via macrophage-mediated efferocytosis. Selective binding of REG3β is achieved by interaction with paucimannosylated proteins that translocate from azurophile granules to the plasma membrane in an activation- and age-dependent manner. Endocytotic uptake and accumulation of REG3β in lysosomes initiates programmed cell death of neutrophils via lyosomal membrane permeabilization and release of cathepsins. Our work establishes REG3β as a local immune checkpoint essential for neutrophil resolution and cardiac repair.

## Main

The myocardium undergoes a series of wound healing processes in response to damage, which eventually result in formation of scars and cardiac remodeling. Immediately following destruction of myocardial tissue different waves of immune cells arrive, a mandatory prerequisite for cardiac repair^1^. The first type of immune cells that infiltrate the infarcted heart in higher numbers are circulating neutrophil granulocytes^2,3^. Neutrophils are recruited to sites of injury for removal of cellular debris but also for release of tissue-degrading enzymes, cytokines, and reactive oxygen species (ROS), which are parts of the inflammatory response^4,5^. On the other hand, neutrophils possess anti-inflammatory and proangiogenic properties that facilitate cardiac repair^6,7^. Despite these beneficial effects, accumulation or belated removal of neutrophils have deleterious consequences, often leading to cardiac rupture^8^. Likewise, elevated neutrophil numbers in the blood are associated with worsened prognosis and increased mortality after MI^9^. The seemingly conflicting functions of neutrophils when coping with consequences of MI may be explained by differential activities and dynamics of different neutrophil subsets. For example, aged neutrophils represent an exceedingly active subset, promoting inflammatory conditions. Tight control of infiltration, maintenance, and persistence of different subsets of neutrophils appears to be crucial for effective wound healing after MI and for avoiding maladaptive remodeling of the heart^10^.

The mechanisms regulating recruitment of neutrophils in response to MI are complex but well characterized^11,12^. In contrast, less is known about local circuits controlling maintenance and removal of different subpopulations of neutrophils after MI. It is assumed that clearance of neutrophils from infarcted hearts is primarily initiated by apoptosis of neutrophils, followed by macrophage-mediated efferocytosis^13^. Activation of the extrinsic apoptotic pathway in neutrophils has been claimed to rely on cell death receptors, namely Fas protein (also called CD95 or APO-1) and receptors for TNFα^14^. However, experimental evidence about the importance and specificity of individual ligands, critical for controlling the life time of different types of neutrophils in the infarcted heart, is mostly missing. In addition to apoptosis, other types of cell death have been described to contribute to the removal of neutrophils in different tissues, such as necroptosis, pyroptosis, necrosis, and NETosis^14^.

We previously described that inactivation of Regenerating-Islet derived protein 3 beta (REG3β) attenuates recruitment of monocytes to the infarcted heart, presumably prolonging persistence of neutrophils. Genetic inactivation of Reg3β frequently causes cardiac rupture and decreases cardiac function in surviving animals after MI^8^. REG3β belongs to the C-type lectin superfamily and is secreted by cardiomyocytes in the border zone of infarcts. The receptors for REG3β or other REGs are essentially unknown. Claims have been made for different membrane proteins to act as receptors for individual REGs, but conclusive evidence is missing and no direct link to intracellular processes downstream of REGs exists^15^. In this study, we discovered that REG3β has direct cytotoxic activity on neutrophils in the infarcted heart, primarily targeting aged neutrophils, which is instrumental for spatial and temporal control of neutrophil presence. REG3β exerts its function by binding to distinct paucimannosidic carbohydrate motifs on the cellular surface of aged and hyperactive neutrophils. Bound REG3β enters neutrophils via endocytosis and accumulates in lysosomes, causing lysosomal membrane permeabilization. Subsequent release of cathepsins and exposure of phosphatidylserine enable clearance of neutrophils via macrophage-mediated efferocytosis.

## Results

### *Reg3b* limits persistence and spatial expansion of neutrophils in the heart after MI

Neutrophils were profiled and quantified in infarcted hearts of wildtype (WT) and *Reg3b* deficient (*Reg3b^-/-^*) mice by multicolor flow cytometry at different time points after permanent ligation of the left anterior descending (LAD) coronary artery. We observed comparable numbers of CD45^hi^/CD11b^hi^/Ly6G^hi^ cardiac tissue neutrophils in WT and *Reg3b^-/-^*mice at day 1 and day 2, suggesting that recruitment of neutrophils does not depend on *Reg3b* (Fig. 1a). In contrast, infarcted hearts of *Reg3b^-/-^* mice contained significantly more neutrophils at day 3 and day 4 compared to control mice. Neutrophils substantially declined in WT mice between days 3 and 4, whereas the reduction of neutrophils in infarcted hearts of *Reg3b^-/-^* mice was moderate (Fig. 1a). No differences in the concentration of neutrophils were detected in the blood of WT and *Reg3b^-/-^*mice at day 4 after MI, indicating that the reason for neutrophil accumulation in *Reg3b^-/-^* hearts lays within and not outside the organ (Fig. 1b). Additional live/dead cell stain analysis identified increased ratios of viable 7-Aminoactinomycin D (7AAD)^-^/Annexin V (AnnV)^-^ neutrophils in *Reg3b^-/-^* mice hearts, accompanied by a decrease of dead 7AAD^-^/AnnV^+^ neutrophils (Fig. 1c).

**Figure 1:**
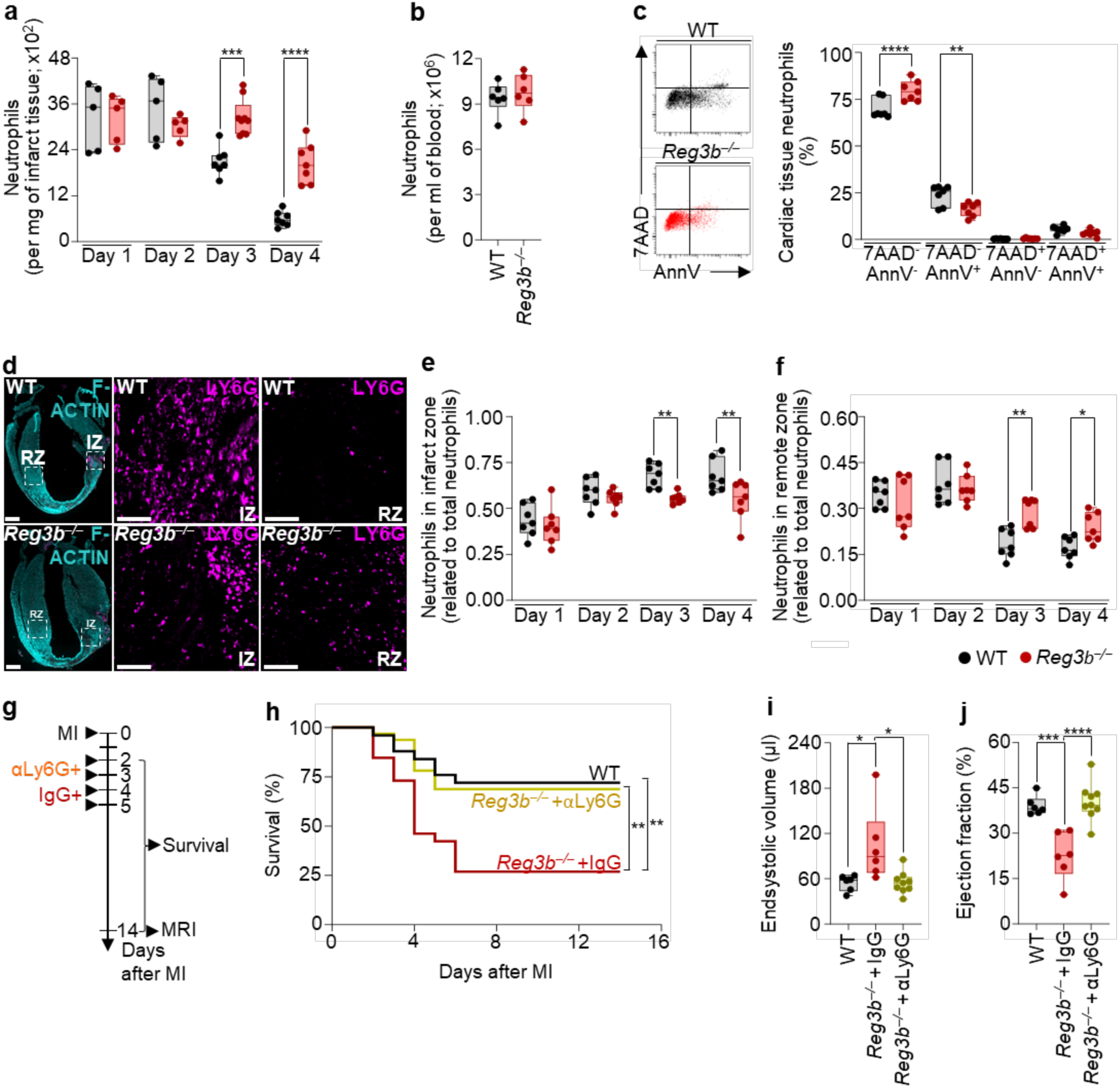
Compromised removal of neutrophils in *Reg3b*-deficient mice after MI increases mortality and impairs heart function. **a**, Flow cytometry quantification of neutrophils in wild-type (WT) and *Reg3b* deficient (*Reg3b^-/-^)* hearts 1, 2, 3, and 4 days after MI. Each data point represents an individual mouse. **b**, Flow cytometry quantification of blood neutrophils in WT and *Reg3b^-/-^*mice 4 days after MI. n = 6 for both groups **c**, Representative flow cytometry dot plots and quantification of viable and dead neutrophils in WT and *Reg3b^-/-^*hearts 4 days after MI, identified by 7-Aminoactinomycin D (7AAD) and Annexin V (AnnV) staining. n = 7 for both groups **d**, **e**, **f**, Immunofluorescent images of LY6G^+^ neutrophils (magenta) in longitudinal sections and zoom-in view of infarcted zone (IZ) and non-infarcted remote zone (RZ) of WT and *Reg3b^-/-^* hearts 4 days after MI. **e**, **f**, Immunofluorescent quantification of Ly6G^+^ neutrophils in serial longitudinal sections of WT and *Reg3b^-/-^*hearts 1, 2, 3, and 4 days after MI. n = 7 mice per group and timepoint. Phalloidin staining was used to distinguish infarcted from non-infarcted tissue. Scale bars, 1000μm and 100μm in magnified sections. **g**, Schematic outline of treatment with control (IgG) or anti-Ly6G (αLy6G) antibodies after MI. **h**, Kaplan-Meier survival curves of WT (n = 26), *Reg3b^-/-^* +IgG (n = 25), and *Reg3b^-/-^* +αLy6G (n = 31) after MI. **i**, **j**, MRI-based analysis of end-systolic volume, and ejection fraction of WT (n = 6), *Reg3b^-/-^* +IgG (n = 6), and *Reg3b^-/-^* +αLy6G (n = 9) mice 14 days after MI. Data are mean ± s.e.m. Two-way ANOVA followed by Sidak’s multiple comparison test (**a**), one-way ANOVA followed by Sidak’s multiple comparison test (**c, i**, **j**), two-way ANOVA followed by Tukey’s multiple comparison test (**e**, **f**), Kaplan-Meier survival analysis (**h**). All experiments were conducted with male mice.

To analyze whether REG3β not only regulates the temporal persistence of neutrophils after MI but also their spatial distribution we performed immunofluorescence staining. Labeling of LY6G^+^ neutrophils uncovered a pronounced expansion of neutrophils from the infarct (IZ) to the left ventricular remote zone (RZ) in *Reg3b^-/-^* hearts at day 3 and 4 after MI (Fig. 1d–f; Suppl. Fig. 1). Fractionation of infarcted hearts into IZ and RZ followed by western blot analysis detected increased concentration of neutrophil-derived proteases including Myeloperoxidase (MPO), Neutrophil elastase (ELANE), and Matrix metalloproteinase-9 (MMP-9) in the RZ at day 4 after infarct, corroborating enhanced spatial expansion of neutrophils in *Reg3b^-/-^* hearts (Suppl. Fig. 2a, b).

We reasoned that the enhanced levels of neutrophil-derived proteases might be responsible for the increased incidence of cardiac rupture in *Reg3b^-/-^* mice, which occurs at the intersection between injured and healthy tissue^8,16^ (Suppl. Fig. 2c). To validate this hypothesis, we transiently depleted neutrophils by injections of anti-LY6G lytic antibodies into *Reg3b^-/-^* mice at day 2, 3, 4, and 5 after MI (Fig. 1g). Anti-LY6G injections efficiently prevented cardiac rupture in *Reg3b^-/-^*mice after MI and increased survival to WT levels (Fig. 1h). Moreover, magnetic resonance imaging-based analysis of heart function revealed that antibody-mediated depletion of neutrophils decreased endsystolic volume and increased ejection fraction of *Reg3b^-/-^* mice 14 days after MI (Fig. 1i, j). The normalization of heart function by injection of anti-LY6G antibodies demonstrates that accumulation of neutrophils is responsible for the impaired function of *Reg3b^-/-^* hearts after MI.

### REG3β directly interacts with neutrophils and initiates cell death

REG3β may affect maintenance of neutrophils either indirectly or by direct interactions. Three-dimensional visualization of LY6G^+^ neutrophils and REG3β protein in mouse hearts after MI via light sheet fluorescence microscopy demonstrated a strong overlap of signals, particularly at the intersection between IZ and RZ (Fig. 2a). Colocalization analysis based on 3D light sheet fluorescence data for LY6G and REG3β in whole infarcted hearts yielded a Pearson’s correlation coefficient of 0.55 (Fig. 2b). Localization of REG3β and the human orthologue REG3A on neutrophils (LY6G^+^ neutrophils in mice and CD66b^+^ neutrophils in humans) was also confirmed by high-resolution fluorescence imaging of infarcted mouse and heart samples from human patients diagnosed with MI (Fig. 2c, d; Suppl. Fig. 3a, b). Flow cytometry analysis uncovered an increase of REG3β-binding (REG3β^pos^) neutrophils in the heart after MI, whereas the numbers of REG3β^pos^ neutrophils in the bone marrow and blood were much lower and not affected by MI (Fig. 2e, f). Notably, we observed comparable ratios of REG3β^pos^ neutrophils in cardiac tissue samples in humans and mice (approximately 11% in human and 15% in mice) (Fig. 2d, e).

**Figure 2:**
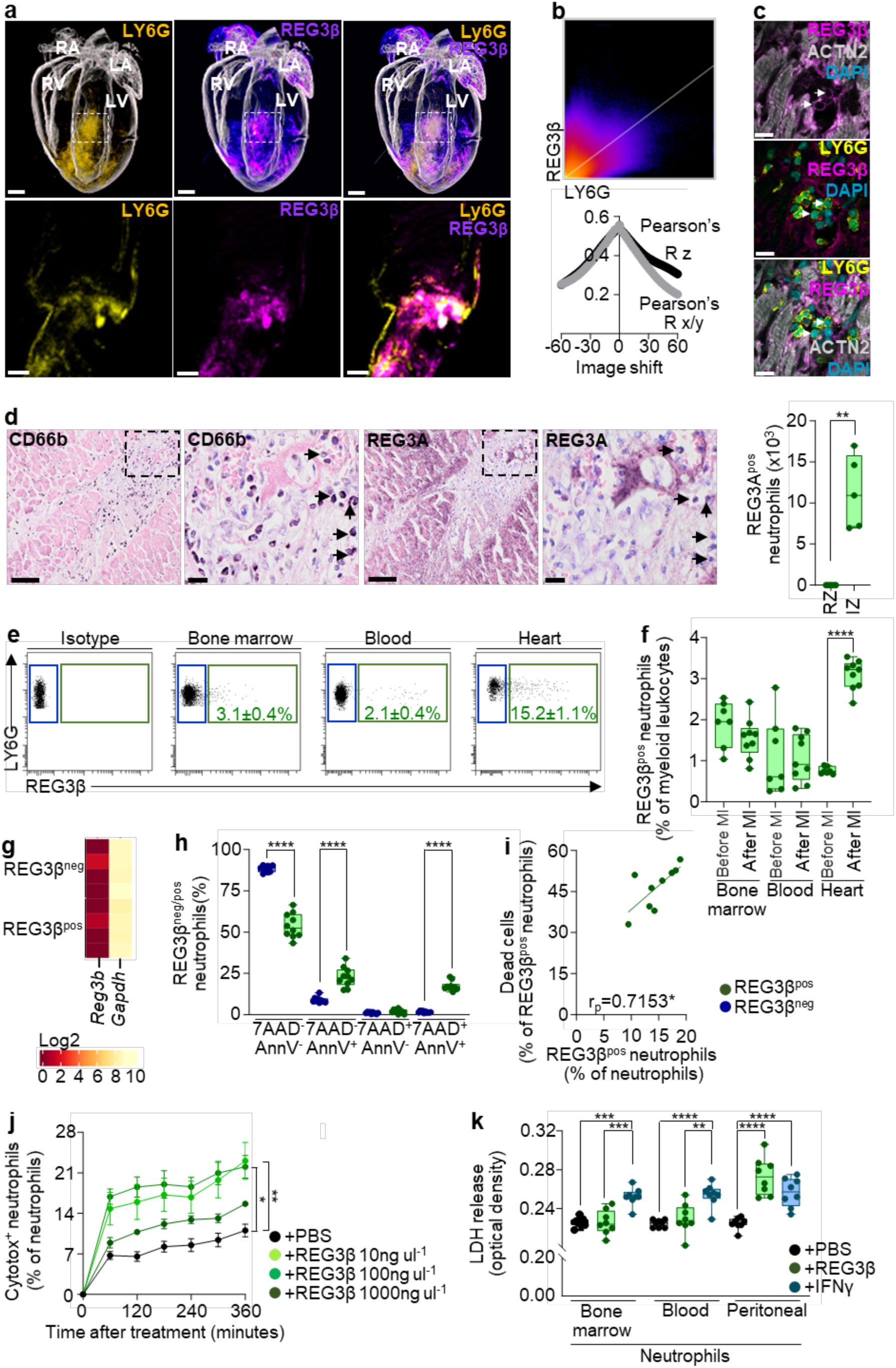
REG3β directly binds neutrophils and initiates rapid cell death. **a**, Representative 3D visualization of LY6G^+^ neutrophils (yellow) and REG3β protein localization (magenta) by light sheet fluorescence microscopy within whole hearts and zoom-in view of WT mice at day 2 after MI. Muscle autofluorescence is white. Left atrium (LA), right atrium (RA), left ventricle (LV), and right ventricle (RV). Scale bars, 1000μm and 500μm in magnified sections. **b**, Co-localization analysis of LY6G and REG3β signal intensities and calculation of correlation coefficient by linear regression. n = 3. **c**, Immunofluorescent images of LY6G^+^ neutrophils (yellow) and REG3β (magenta) protein localization in hearts of WT mice 2 days after MI. Alpha-actinin-2 (ACTN2): grey and 4′,6-Diamidin-2-phenylindol (DAPI): blue. Scale bars, 10μm. **d**, Immunohistochemical 3,3’-Diaminobenzidine (DAB) visualization of CD66b^+^ neutrophils and REG3A in myocardial biopsies of humans with MI. Arrows indicate REG3A^+^ neutrophils. Counter staining with hematoxylin. Scale bars, 100μm. **e**, Representative flow cytometric dot plots of REG3β-negative (REG3β^neg^, blue) and REG3β-positive (REG3β^pos^, green) neutrophil subsets from bone marrow, blood and heart of WT mice 2 days after MI. Mean ± sem of REG3β^pos^ neutrophils in % of neutrophils are shown in each dot plot. n = 10. **f**, Flow cytometric quantification of REG3β^pos^ neutrophils in % of myeloid leukocytes in bone marrow, blood and heart of WT mice before (n = 7) and 2 days after MI (n = 9). **g**, Heatmap of *Reg3b* gene expression in REG3β^neg^ and REG3β^pos^ neutrophils from cardiac tissue of WT mice 2 days after infarct. *Gapdh* was used as reference. n = 4. **h**, Flow cytometric quantification of viable and dead REG3β^neg^ and REG3β^pos^ neutrophils by Annexin V and 7AAD staining in WT mice 2 days after MI. n = 10. **i**, Pearson correlation analysis between REG3β^pos^ and cell death. n = 9. **j**, Cell death kinetics of peritoneal neutrophils from WT mice, treated with REG3β at indicated concentrations, determined by incorporation of Cytotox. Treatment with PBS served as control. n = 7 for all groups. **k**, Lactate dehydrogenase (LDH) release of isolated bone marrow, blood and peritoneal neutrophils from WT mice after treatment with REG3β (100ng ul^-1^). Administration of PBS and interferon gamma (IFNγ, 100ng ml^-1^) was used as controls. N = 8 for all groups. Data are mean ± s.e.m. Pearson correlation (**b**, **i**), nonparametric Kolmogorov-Smirnov test (**d**), one-way ANOVA followed by Sidak’s multiple comparison test (**f**, **h**), two-way ANOVA mixed-effects analysis followed by Sidak’s multiple comparison test (**j**), two-way ANOVA followed by Tukey’s multiple comparison test (**k**). All experiments were conducted with male mice.

We previously described that the main source of REG3β in the infarcted heart are cardiomyocytes^8,17^. Nevertheless, we wanted to make sure that the REG3β signal on neutrophils is indeed derived from secreted REG3β and not from REG3β translated within neutrophils. Bulk RNA-sequencing of sorted REG3β^neg^ and REG3β^pos^ neutrophils from infarcted hearts did not detect any transcripts of *Reg3b* in either population, corroborating previous data (Fig. 2g). Interestingly, the ratio of neutrophils with intracellular localization of REG3β was substantially higher than the ratio of neutrophils with REG3β at the cell surface, suggesting that bound REG3β is rapidly taken up by neutrophils (Suppl. Fig. 4a, b). Even more importantly, REG3β^pos^ neutrophils, either intracellular or membrane bound, showed reduced viability (Fig. 2h, I; Suppl. Fig. 4c).

The observation that REG3β^pos^ neutrophils showed reduced viability prompted us to test potential direct cytotoxic effects of REG3β on neutrophils. We isolated mouse neutrophils from peritoneal cavities after induction of peritonitis with casein, treated them with different concentrations of REG3β, and monitored cell death with the IncuCyte Cytotox Green Assay in real time. We observed a rapid increase of dead Cytotox^+^ neutrophils already after 60 minutes. Potent cytotoxic effects were recorded at 10 ng ml^-1^ and 100ng ml^-1^ REG3β, whereas cytotoxicity declined at 1000ng ml^-1^, probably due to enhanced aggregation of REG3β (Fig. 2j). Lactate dehydrogenase (LDH) release assays confirmed cytotoxic effects of recombinant REG3β protein on activated neutrophils. In stark contrast, bone marrow- and blood-derived neutrophils, which are in a more quiescent and non-inflammatory state, were not responding to REG3β (Fig. 2k)^18^. Importantly, interferon gamma (IFNγ), which was used as a positive control did not show any selectivity in respect to activated or non-inflammatory states but exerted cytotoxic effects on all types of neutrophils (Fig. 2k).

### REG3β binds to hyperactive and aged neutrophils

Our data indicated that REG3β did not bind to all but a subset of neutrophils. To characterize REG3β^pos^ neutrophils more closely, we performed RNA sequence analysis of REG3β^neg^ and REG3β^pos^ neutrophils from infarcted hearts. Subsequent principal component analysis separated REG3β^neg^ and REG3β^pos^ neutrophils into two distinct subsets (Fig. 3a). Compared to REG3β^neg^ neutrophils 1124 genes were up- and 285 genes were downregulated in REG3β^pos^ neutrophils (Fig. 3b).

**Figure 3:**
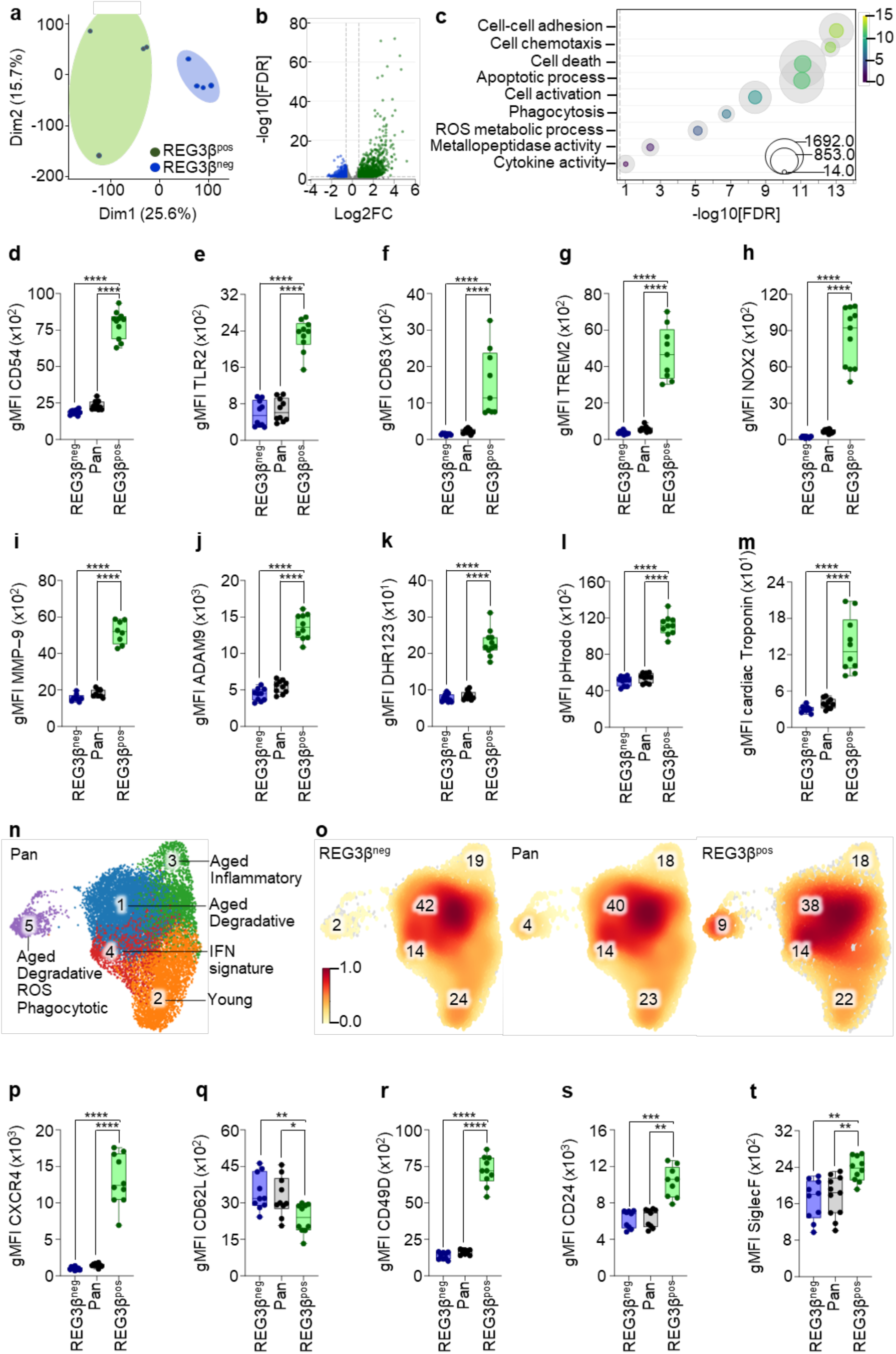
REG3β binds to hyperactive, aged neutrophils in the infarcted heart. **a**, Principal component analysis of REG3β-negative (REG3β^neg^, blue) and REG3β-positive (REG3β^pos^, green) neutrophils obtained from WT hearts 2 days after MI. (n = 4 for both groups). **b**, Volcano plot depicting fold changes (FC) and False Discovery Rate (FDR)-adjusted p-values of differentially expressed genes (DEG) in REG3β^neg^ and REG3β^pos^ cardiac tissue neutrophils. **c**, KOBAS (KEGG Orthology-Based Annotation System) pathway enrichment bubble plot of DEGs of REG3β^neg^ compared with REG3β^pos^ heart neutrophils. Selected significantly enriched pathways are shown (*P*<0.05). The dashed line marks the *P* value of 0.05. Outer gray circles represent the total number of genes in each pathway. Centered colored circles represent the number of DEG in each pathway. Diameter and color of circles indicate the number of genes and significance of enrichment, respectively. **d–h**, Geometric mean fluorescence intensity (gMFI) of CD54 (**d,** n = 11), TLR2 (**e,** n = 10), CD63 (**f**, n = 9), TREM2 (**g**, n = 9), and NOX2 (**h**, n = 11) on REG3β^neg^, Pan and REG3β^pos^ neutrophils from WT hearts 2 days after MI. **i–m**, gMFI of MMP-9 (**i**, n = 8), and ADAM9 (**j**, n = 10), DHR123 (**k**, n = 11), pHrodo (**l**, n = 10), and cardiac troponin (**m**, n = 10) in REG3β^neg^, pan, and REG3β^pos^ neutrophils from WT hearts 2 days after MI. **n**, Uniform manifold approximation and projection (UMAP) of neutrophil gene expression obtained from WT hearts at day 2 after MI. **o**, Density plots of REG3β^neg^, pan and REG3β^pos^ neutrophils from WT hearts 2 days after MI. Cluster ratios for REG3β^neg^, pan and REG3β^pos^ heart neutrophils are also shown. **p**–**t**, gMFI of CXCR4 (**p,** n = 10), CD62L (**q,** n = 10), CD49D (**r**, n = 10), CD24 (**s**, n = 9), and SiglecF (**t**, n = 10) on REG3β^neg^, pan and REG3β^pos^neutrophils from WT hearts 2 days after MI. Data are mean ± s.e.m. One-way ANOVA followed by Sidak’s multiple comparison test (**d**-**m**, **p**-**r, t**), Kruskal-Wallis 1-way ANOVA followed by Dunn’s multiple comparison test (**s**). All experiments were conducted with male mice.

Selective pathway enrichment analysis using KOBAS (KEGG Orthology-Based Annotation System) demonstrated increased expression of genes related to cell death in REG3β^pos^ neutrophils. Furthermore, we detected increased expression of markers for nearly all critical neutrophil effector functions, including cell adhesion, chemotaxis, activation, phagocytosis, ROS metabolism, metallopeptidase and cytokine activity (Fig. 3c). Flow cytometric analysis corroborated enhanced presence of neutrophil activation markers such as CD54, TLR2, CD63, and TREM2 in REG3β^pos^ compared to pan and REG3β^neg^ neutrophils (Fig. 3d–g). Levels of the ROS producing enzyme NOX2 and the metallopeptidases MMP-9 and ADAM9 were substantially elevated in REG3β^pos^ neutrophils (Fig. 3h–j). We also measured increased production of ROS with DHR123 and increased phagocytotic activity by incorporation of pHrodo and cardiomyocyte-derived cardiac troponin (Fig. 3k–m). The concentration of REG3β^pos^ neutrophils was much lower in the bone marrow and blood compared to the infarcted heart but, if present, showed the same hyperactivated state, indicated by increased geometric mean fluorescent intensities (gMFI) of CD54, CD63, NOX2 and DHR123 compared with REG3β^neg^ neutrophils (Suppl. Fig. 5a–d).

To position REG3β^pos^ neutrophils in the continuum of neutrophil activation and maturation, we subjected REG3β^neg^ and REG3β^pos^ neutrophils sorted from infarcted hearts to scRNA-seq^19,20^. Analysis of 5180 REG3β^pos^ and 10814 REG3β^neg^ neutrophils with a median of 1248 expressed genes per cell identified five different subsets of neutrophils in the infarcted mouse heart (Fig. 3n; Suppl. Fig. 6a). Next, we calculated bone marrow proximity (BMP) and aging scores of each cluster according to studies analyzing neutrophil development and aging^21^. Cluster 2 showed the highest BMP score with gene expression characteristic for young neutrophils (*Sell*, *Retnlg*, *Lcn2*), whereas clusters 1, 3, and 5 had higher aging scores and expressed genes typical for aged neutrophils (*Cxcr4*, *Siglecf*, *Ncf1*) (Supp. Fig. 6b–e). Pseudotime analysis, defining cluster 2 as root, placed clusters 1, 3, and 5 at the end of the neutrophil maturation trajectory (Suppl. Fig. 6f). We also identified a unique differentiated subset of neutrophils, cluster 4, characterized by increased expression of interferon (IFN) stimulated genes (*Irf7*, *Ifit1*, *Ifi204*), whose presence in the heart was independent of MI and did not follow the neutrophil differentiation trajectory (Suppl. Fig. 6g, h)^19,22^.

Most REG3β^pos^ neutrophils belonged to the clusters of aged neutrophils, which was confirmed by flow cytometric analysis, showing increased presence of neutrophil aging markers such as CXR4, CD49D, CD24, and SiglecF in as REG3β^pos^ neutrophils and decreased presence of CD62L, a marker for young neutrophils (Fig.3 o–t)^19^. A particular enrichment of REG3β^pos^ neutrophils was seen in aged cluster 5 (9% of REG3β^pos^ vs. 2% of REG3β^neg^), which has the highest scores for metallopeptidase activity, ROS production, and phagocytosis among all neutrophil subsets (Fig. 3n, o; Suppl. Fig. 6i–p). To analyze whether activation and aging is a prerequisite for binding and subsequent cytotoxicity of REG3β or a mere epiphenomenon, we cultured isolated neutrophils from the bone marrow and exposed them to lipopolysaccharide (LPS) to induce activation and aging (Suppl. Fig. 7a–d). As expected, activation of bone marrow-derived neutrophils increased REG3β binding and REG3β-dependent cytotoxicity (Suppl. Fig. 7e–i). Taken together, the data demonstrate that REG3β preferentially binds and kills hyperactivated, aged neutrophils.

### Translocation of granule-derived paucimannose-conjugated proteins to the cell surface enables binding of REG3β

Since no conclusive evidence of a receptor for REG3β exists but specific interactions of REG proteins with carbohydrate epitopes of peptidoglycans on the cellular surface of bacterial cells have been reported^15,23,24^, we reasoned that REG3β may exerts its action via binding to cell surface N-glycans. Cell surface N-glycan analysis of sorted REG3β^neg^ and REG3β^pos^ peritoneal neutrophils identified 28 N-glycans, which were grouped into 8 distinct glycan traits (Suppl. Fig. 8a–c; Suppl. Table 2). High mannose structures dominated the cell surface N-glycome of REG3β^neg^ and REG3β^pos^ neutrophils, followed by presence of fucose, galactose, and complex N-glycan traits on REG3β^neg^ neutrophils (Suppl. Fig. 8c). In contrast, 18% of the cell surface N-glycome of REG3β^pos^ but only 2% of REG3β^neg^ neutrophils consisted of atypical small paucimannose type N-glycans, which is an under-studied class of N-glycosylation in mammalian cells (Fig. 4a; Suppl. Fig. 8c)^25,26,27^. Immunofluorescence staining and flow cytometric analysis using the paucimannose-reactive Mannitou antibody confirmed enrichment for paucimannosylation of REG3β^pos^ neutrophils^28^ (Fig. 4b, c).

**Figure 4:**
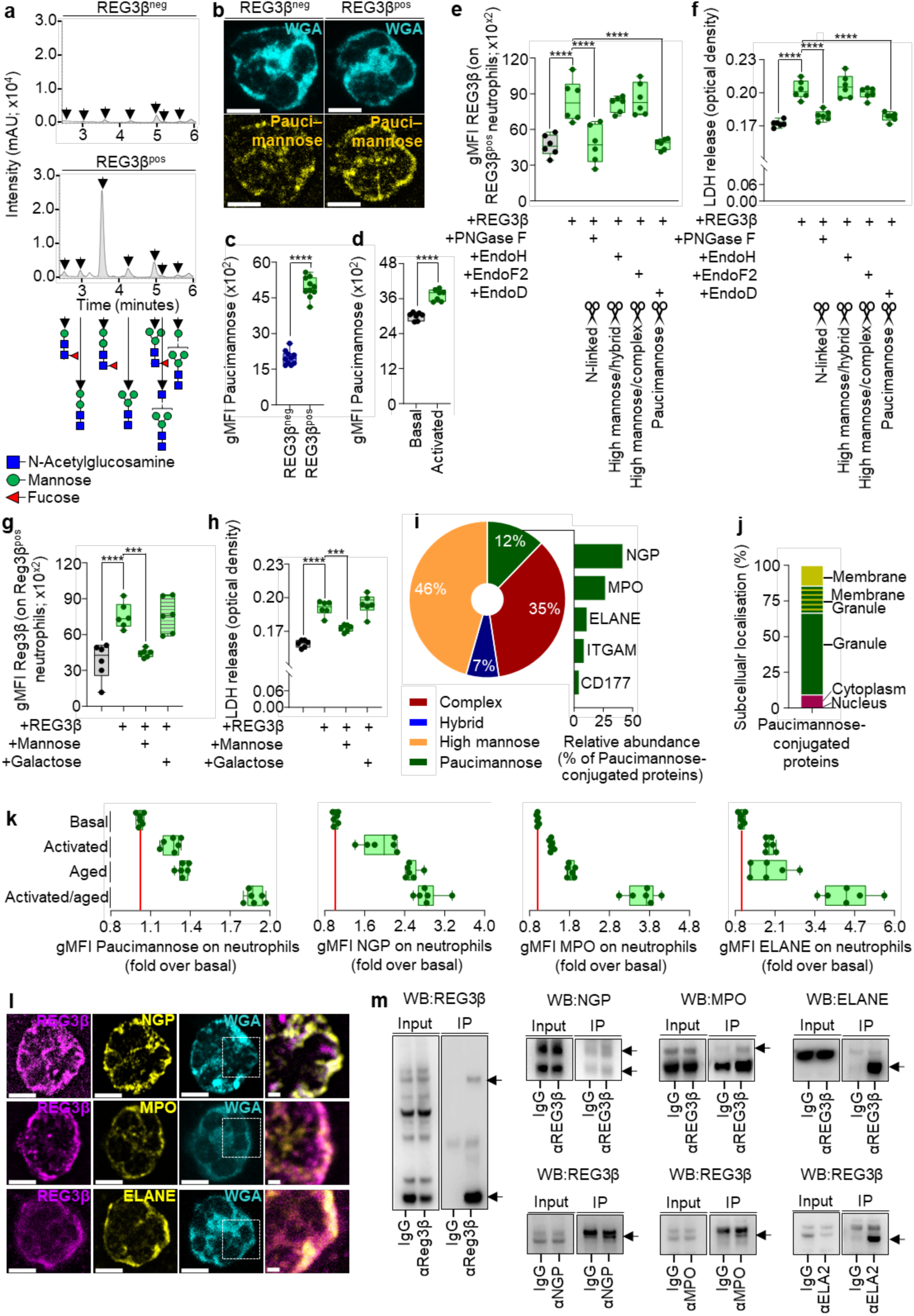
Binding of REG3β to neutrophil subsets is enabled by translocation of granule-derived paucimannose-conjugated proteins. **a**, Cell surface N-glycan analysis of sorted REG3β-negative (REG3β^neg^) and REG3β-positive (REG3β^pos^) peritoneal neutrophils from WT mice. Zoomed-in view chromatograms of Paucimannose-type N-glycans from REG3β^neg^ and REG3β^pos^ neutrophils are shown. mAU, milliabsorbance units. **b**, Immunofluorescent images of paucimannose (yellow) in REG3β^neg^ and REG3β^pos^ peritoneal neutrophils from WT mice. Wheat germ agglutinin (WGA): grey. Scale bars, 5μm. **c**, Geometric mean fluorescence intensity (gMFI) of paucimannose on REG3β^neg^ and REG3β^pos^ cardiac tissue neutrophils from WT mice 2 days after MI. n = 10. **d**, gMFI of Paucimannose on basal versus activated bone marrow neutrophils. n = 7. **e**, gMFI of REG3β on REG3β^pos^ bone marrow derived neutrophils pretreated with peptide:N-glycosidase F (PNGase F), endoglycosidases H (EndoH), F2 (EndoF2), and D (EndoD) (all used at 1U ml^-1^) for 30 minutes after stimulation with REG3β (100 ng ml^-1^) for 15 minutes. PBS served as control. n = 6. **f**, Lactate dehydrogenase (LDH) release of activated bone marrow derived-neutrophils pretreated with PNGase F, EndoH, EndoF2, and EndoD (all used at 1U ml^-1^) for 30 minutes following stimulation with REG3β (100 ng ml^-1^) for 30 minutes. Cleavage specificity of each glycosidase is indicated. PBS served as control. n = 6. **g**, **h**, gMFI of REG3β on REG3β^pos^ bone marrow-derived neutrophils (**g**, n = 6) and LDH release of activated bone marrow derived-neutrophils (**h**, n = 6) treated with Mannose, Galactose (both 100 mM) and REG3β (100 ng ml^-1^) for 30 minutes. PBS served as control. **i**, Relative abundance of major N-glycan groups in peritoneal neutrophils. Relative abundance of the five most abundant Paucimannose-conjugated proteins including neutrophilic granule protein (NGP), myeloperoxidase (MPO), neutrophil elastase (ELANE), integrin alpha M (ITGAM) and CD177 antigen (CD177). **j**, Subcellular localization of paucimannose-conjugated proteins in peritoneal neutrophils. **k**, Ratio of gMFI of paucimannose, NGP, MPO, and ELANE on basal, activated, aged, and activated/aged bone marrow derived-neutrophils in relation to basal-state neutrophils. Ratios are shown as fold change, whereas levels in basal cells is set to 1 and illustrated by a red line. n = 6. **l**, Immunofluorescent images of NGP, MPO, and ELANE (all in yellow) in non-permeabilized REG3β^pos^ peritoneal neutrophils from WT mice. REG3β: magenta and WGA: grey. Scale bars, 5μm and 1μm in magnified sections. **m**, Immunoblot analysis of input and cell surface co-immunoprecipitated samples from peritoneal neutrophils using antibodies against REG3β, NGP, MPO and ELANE. Isotype controls (IgG) were used as controls. Arrows indicate bands co-precipitated with the primary antigen. Data are mean ± s.e.m. Two-sided unpaired t tests (**c, d**), one-way ANOVA followed by Sidak’s multiple comparison test (**d**–**g**). All experiments were conducted with male mice.

Furthermore, activation of bone-marrow derived neutrophils with LPS elevated paucimannose levels on the cell surface, suggesting that the ability of REG3β to bind to activated neutrophils is mediated by paucimannosylation (Fig. 4d; Suppl. Fig. 7e). To test this hypothesis, we treated activated neutrophils with the enzyme Peptide:N-glycosidase F (PNGase F), which cleaves all N-linked glycans from glycoproteins. PNGase F treatment essentially abolished binding of REG3β to activated neutrophils (Fig. 4e) and also neutralized cytotoxic effects (Fig. 4e, f). Importantly, treatment of activated neutrophils with endoglycosidase D (EndoD), specifically cleaving Paucimannose type N-glycans, caused similar effects (Fig. 4e, f), whereas treatment with Endo H or EndoF2, which cleave high mannose hybrid and complex saccharides, respectively, did neither affect REG3β binding nor cytotoxicity (Fig. 4e, f). To further corroborate these results, we performed competition assays with different monosaccharides in millimolar concentrations. We found that mannose, a key component of paucimannosidic N-glycans, but not galactose inhibited binding of REG3β and prevented cell death (Fig. 4f, g).

To identify paucimannosylated glycoproteins interacting with REG3β, we conducted a liquid chromatography tandem mass spectrometry (LC-MS/MS)-based glycoproteomic analysis of peritoneal neutrophils. We found that the majority of glycoproteins were primarily decorated with high mannose, followed by complex, paucimannose and hybrid glycans (Fig. 4h). In total, 21 paucimannosylated proteins were identified. Mannose_2_Fucose_1_N-Acetylglucosamine_2_ (M2F) was the most abundant paucimannose signature of REG3β^pos^ neutrophils, a characteristic N-glycan of azurophile granule derived glycoproteins (Fig. 6a; Suppl. Fig. 9b)^26^. The vast majority of paucimannosylated proteins (approx. 94%) consisted of neutrophilic granule protein (NGP), myeloperoxidase (MPO), neutrophil elastase (ELANE), integrin alpha M (ITGAM), and CD177 (CD177) (Fig. 4h; Suppl. Fig. 9a, b). Analysis of the putative subcellular localization by neXtprot indicated that NGP, MPO, and ELANE reside in azurophilic granules (Fig. 4i; Suppl. Fig. 9c)^25,27^. Based on these results we reasoned that paucimannosylated granular proteins of REG3β^pos^ neutrophils are translocated to the cell membrane following activation and subsequent degranulation of neutrophils.

Flow cytometric cell surface profiling for paucimannosylated proteins confirmed our hypothesis, revealing elevated levels of NGP, MPO, and ELANE on the cell membrane of cardiac tissue REG3β^pos^ neutrophils (Suppl. Fig. 10a–c), whereas ITGAM and CD177 were either not altered or declined (Suppl. Fig. 10d, e). Furthermore, we detected a rapid increase of paucimannose, NGP, MPO, and ELANE upon activation of neutrophils, which further increased with aging and peaked in hyperactivated, aged neutrophils (Fig. 4k; Suppl. Fig. 10f–h). Concomitant increase of extracellular levels of the degranulation marker CD63 support the idea that increased degranulation in activated and aging neutrophils maximizes translocation of paucimannosylated proteins (Suppl. Fig. 10i). Moreover, we found that inhibition of neutrophil degranulation by Nexinhib20 diminished binding of REG3β to activated neutrophils and abrogated cytotoxic effects (Suppl. Fig. 10j, k)^29^. To interrogate membrane-specific interactions of REG3β with azurophile granule derived paucimannosylated proteins, we used confocal microscopy, which revealed the presence of NGP, MPO, and ELANE on wheat germ agglutinin (WGA)^+^ plasma membranes of REG3β^pos^ neutrophils, partially colocalizing with REG3β (Fig. 4l). Cell surface co-immunoprecipitation experiments using REG3β-treated neutrophils uncovered interactions between REG3β and NGP, MPO, and ELANE at the plasma membrane (Fig. 4m; Suppl. Fig. 11 a–d, f–k). No interactions were detected between REG3β and ITGAM and CD177, illustrating specificity of these interactions (Suppl. Fig. 11d, e). Taken together, our results demonstrate that paucimannosylated proteins are redistributed by degranulation to the cell surface of neutrophils in an activation- and age-dependent manner, enabling binding of REG3β.

### REG3β induces lysosome-mediated cell death of activated neutrophils

Induction of apoptosis by tumor necrosis factor alpha (TNFα) is an established mechanism to accomplish programmed cell death of neutrophils, although data for cardiac neutrophils are scarce^30^. To investigate whether REG3β employs a similar machinery to kill neutrophils, we directly compared effects of REG3β and TNFα on neutrophils. We observed similar kinetics of LDH release kinetics and loss of Mitospy, a fluorescent reagent labelling mitochondria in viable cells after treatment with either REG3β or TNFα (Fig. 5a, b). The ratio of 7AAD^-^/AnnV^+^ neutrophils and formation of forward-side-scatter (FSC)^low^/AnnV^+^ small vesicles increased in REG3β- or TNFα-treated neutrophil cultures, though the increase of 7AAD^-^/AnnV^+^ neutrophils by REG3β was less prominent and only became significant after 60 minutes (Fig. 5c, d). Importantly, we did not detect increased activation of the key executioner caspases 3 and 7 upon administration of REG3β to neutrophils at any time point, in stark contrast to the effects of TNFα (Fig. 5e). We concluded that REG3β does not induce classical apoptotic cell death via activation of caspases 3 and 7.

**Figure 5:**
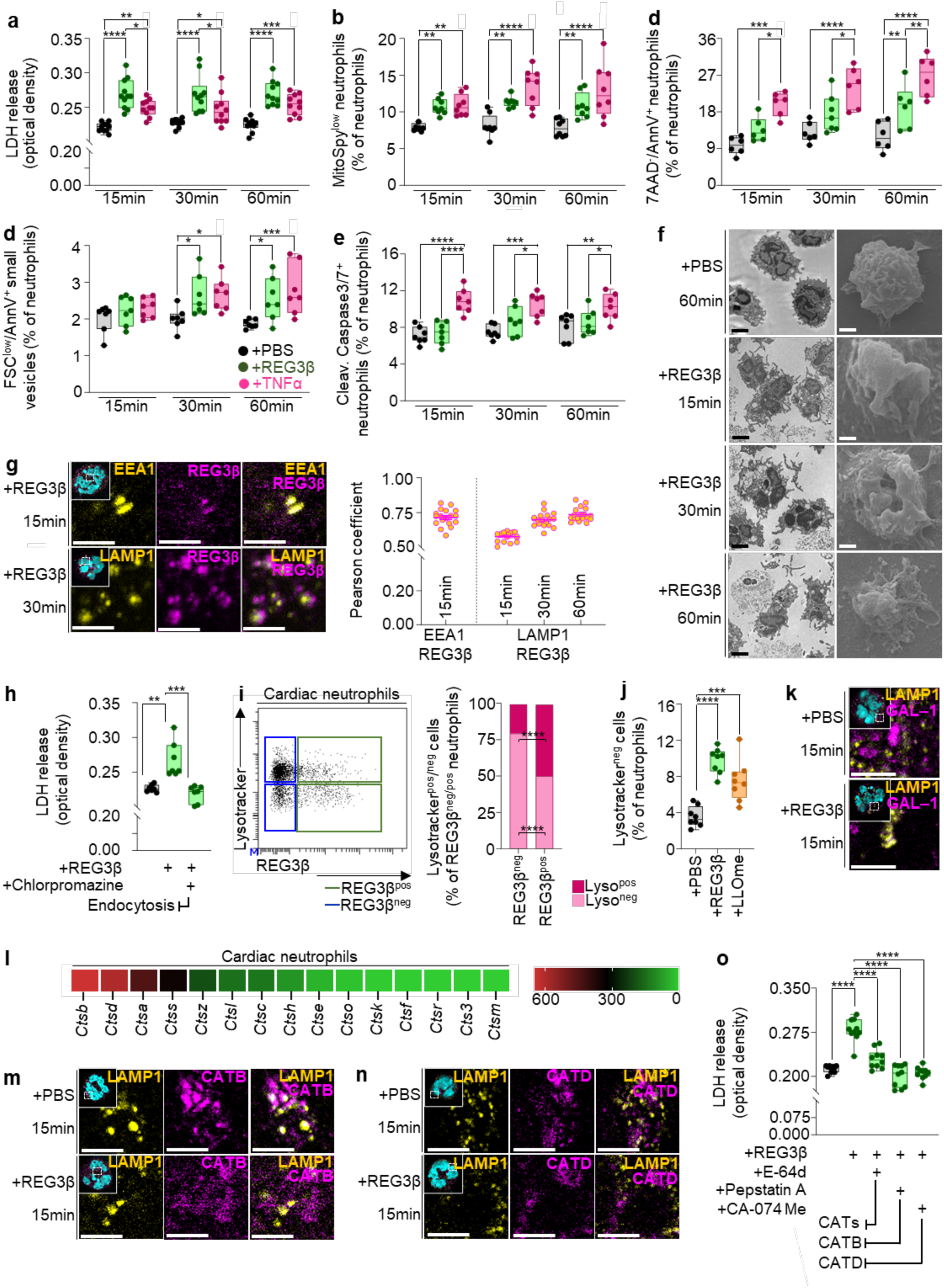
REG3β-induced release of lysosomal cathepsins causes cell death. **a–e**, Lactate dehydrogenase (LDH) release (**a**, n = 10 for all groups), flow cytometric quantification of Mitospy^low^ (**b**, n = 8 for all groups), quantification of 7AAD^-^/AnnV^+^ (**c**, n = 6 for all groups), quantification of FSC^low^/AnnV^+^ small vesicles (**d**, n = 6 for all groups), and quantification of cleaved caspase 3/7 activity (**e**, n = 7 for all groups) of REG3β-treated (100ng ul^-1^) peritoneal neutrophils from WT mice at indicated time points. PBS and tumor necrosis factor alpha (TNFα, 100ng ml^-1^) were used as controls. **f**, Transmission (TEM, left panel) and scanning electron microscopy (SEM, right panel) images of peritoneal neutrophils from WT mice treated with REG3β (100ng ul^-1^) at indicated time points. PBS served as control. Scale bars, 2μm for TEM and 2.5μm for SEM. **g**, Immunofluorescent images of early endosome antigen 1 (EEA1) combined with REG3β (magenta) and lysosomal-associated membrane protein 1 (LAMP1, yellow) combined with REG3β in REG3β-treated (100ng ul^-1^) peritoneal neutrophils from WT mice at indicated time points. Pearson coefficient of colocalization of EEA1 and LAMP1 with REG3β. n = 18. Scale bars, 5μm. **h**, LDH release of peritoneal neutrophils from WT mice pretreated with chlorpromazine (1μM) for 30 minutes, following stimulation with REG3β (100 ng ml^-1^) for 15 minutes. DMSO served as control. n = 7. **i**, Flow cytometric dot plots and quantification of REG3β^neg^ (blue) and REG3β^pos^ (green) neutrophils from WT mice 2 days after MI, separated into Lyso^pos^ and Lyso^neg^ subsets. n = 7. **j**, Flow cytometric quantification of Lyso^neg^ cells of peritoneal neutrophils from WT mice after treatment with Reg3β (100ng ul^-1^) for 15 minutes. Administration of PBS and recombinant L-leucyl-L-leucine methyl ester (LLOme, 100ng ml^-1^) were used as controls. n = 8 for all groups. **k**, Immunofluorescent images of LAMP1 (yellow) and Galectin-1 (GAL-1, magenta) localization in peritoneal neutrophils from WT mice after treatment with Reg3β (100ng ul^-1^) at indicated time points. PBS served as control. Scale bars, 10μm. **l**, Heatmap of mean cathepsin family gene expression counts in neutrophils from WT hearts 2 days after MI. n = 4. **m**, Immunofluorescent images of LAMP1 (yellow) and Cathepsin B (CATB, magenta), and **n**, LAMP1 (yellow) and Cathepsin D (CATD, magenta) localization in peritoneal neutrophils from WT mice upon treatment with recombinant Reg3β (100ng ul^-1^) for 15 minutes. Administration of PBS served as negative control. Scale bars, 5μm. **o**, LDH release of peritoneal neutrophils from WT mice pretreated with Pan cathepsin inhibitor E-64d, CATD inhibitor Pepstatin A, and CATB inhibitor CA-074Me (all used at 1μM) for 30 minutes following stimulation with recombinant REG3β (100 ng ml^-1^) for 15 minutes. DMSO served as control. n = 10. Data are mean ± s.e.m. Two-way ANOVA followed by Tukey’s multiple comparison test (**b**–**f,**), one-way ANOVA followed by Sidak’s multiple comparison test (**h–j, n**). All experiments were conducted with male mice.

To obtain further insights into the mechanism by which REG3β eliminates neutrophils, we analyzed REG3β-treated cultured peritoneal neutrophils by transmission (TEM) and scanning electron microscopy (SEM). We observed several morphological alterations including cytoplasmatic vacuolization, loss of granular structures and membrane ruffling, which became apparent already after 15 minutes and increased within 60 minutes after exposure to REG3β (Fig. 5f). Characteristic features of caspases 3/7-dependent apoptosis such as chromatin compaction in crescent shaped masses at the nuclear periphery or membrane blabbing were not detected. Due to the presence of REG3β in the cytoplasm of neutrophils, we speculated that REG3β is taken up by activated neutrophils after binding to paucimannosylated membrane proteins and induces cell death by an intracellular mechanism. Proteins are frequently taken up via the endocytotic pathway, involving vesicle formation at the plasma membrane followed by fusion with endosomes, which mature into lysosomes or fuse with preexisting lysosomes^31,32^. Immunostaining combined with expansion microscopy detected REG3β in early endosome antigen 1 (EEA1)^+^ endosomes within minutes upon administration as well as in lysosomal-associated membrane protein 1 (LAMP1)^+^ lysosomes (Fig. 5g). A major endocytic pathway in mammalian cells is clathrin-mediated endocytosis (CME)^33^. To investigate its involvement in the uptake of REG3β, we treated neutrophils with chlorpromazine, a CME blocker^34^. Treatment with chlorpromazine abrogated cytotoxic effects of REG3β on neutrophils (Fig. 5h), indicating a critical role of CME for REG3β-dependent cell death.

To analyze putative direct effects of REG3β on lysosomes, we stained REG3β^neg^ and REG3β^pos^ cardiac neutrophils with Lysotracker, a cell-permeable fluorescent dye labelling acidic lysosomes. Interestingly, REG3β-binding neutrophils showed depletion of lysosomes. Approximately 49% of REG3β^pos^ neutrophils were negative for Lysotracker (Lyso^neg^), but only 21% of REG3β^neg^ neutrophils, indicating depletion of lysosomes in REG3β^pos^ neutrophils (Fig. 5i). Next, we treated Lysotracker-stained peritoneal neutrophils with REG3β, which resulted in a rapid increase of Lyso^neg^ neutrophils. The effects of REG3β were similar to L-leucyl-L-leucine methyl ester (LLOme), a dipeptide, which polymerizes inside lysosomes and induces lysosomal membrane damage (Fig. 5j)^35^. To corroborate REG3β-dependent damage of lysosomal membranes, we employed the galectin puncta assay, which is based on translocation of cytosolic galectins to the endolysosomal glycocalyx after lysosomal membrane permeabilization^36^. We observed a punctuated localization of Galectin-1 (GAL-1), overlapping with LAMP1^+^ lysosomes, whereas PBS-treated neutrophils showed diffuse cytosolic GAL-1 expression, characteristic for intact lysosomes (Fig. 5k).

Lysosomal membrane permeabilization leads to leakage of lysosomal hydrolases, especially cathepsins, which induce cell death in various cell types^37,38^. RNA-seq analysis identified 15 members of the large family of cathepsins in cardiac neutrophils after MI, with Cathepsin B (CATB)- and Cathepsin D (CATD) showing the highest expression (Fig. 5l). Expansion microscopy localized CATB and CATD to lysosomal LAMP1^+^ compartments under basal conditions (Fig. 5m, n). In contrast, administration of REG3β caused a much more diffuse positioning of CATB and CATD in neutrophils, indicating release from lysosomes into the cytosol (Fig. 5m, n). To confirm the role of cathepsin-release from lysosomes for REG3β-mediated cell death, we pretreated neutrophils with pan-cathepsin inhibitor E-64d, CATB inhibitor Pepstatin A, and CATD inhibitor CA-074. All three inhibitors prevented REG3β-induced cell death of activated neutrophils as indicated by reduced release of LDH (Fig. 5o). Taken together our data indicate that REG3β induces death of activated neutrophils by lysosomal membrane permeabilization and release of cathepsins after endocytotic uptake and transport to lysosomes.

### REG3β is required for clearance of neutrophils by efferocytosis

Removal of dying neutrophils from sites of injury is eventually achieved by efferocytosis, employing professional phagocytes, including macrophages, and to a lesser extent by monocytes, dendritic cells and neutrophils^39,40^. In addition to increased localization of AnnV on the cell surface of REG3β^pos^ neutrophils, reflecting externalization of the ‘eat-me’ signal phosphatidylserine, we detected increased expression of genes coding for efferocytotic receptors and “bridging” ligands in REG3β^pos^ neutrophils localized in the scRNA-seq cluster 5 (Fig. 2h; Fig. 6a, b). The cluster also contained macrophage-specific genes such as *Ccr2*, *Mrc1*, *Cd209a* suggesting formation of macrophage-Reg3β^pos^ neutrophil-hybrids. Morphological analysis of sorted cardiac REG3β^neg^ versus REG3β^pos^ neutrophils confirmed this assumption (Fig. 6c, d). Flow cytometry analysis revealed that approx. 60% of REG3β^pos^ neutrophils formed hybrids with CD64^hi^/MERTK^hi^ macrophages, whereas such hybrids were essentially absent when analyzing REG3β^neg^ neutrophils (Fig. 6e).

**Figure 6:**
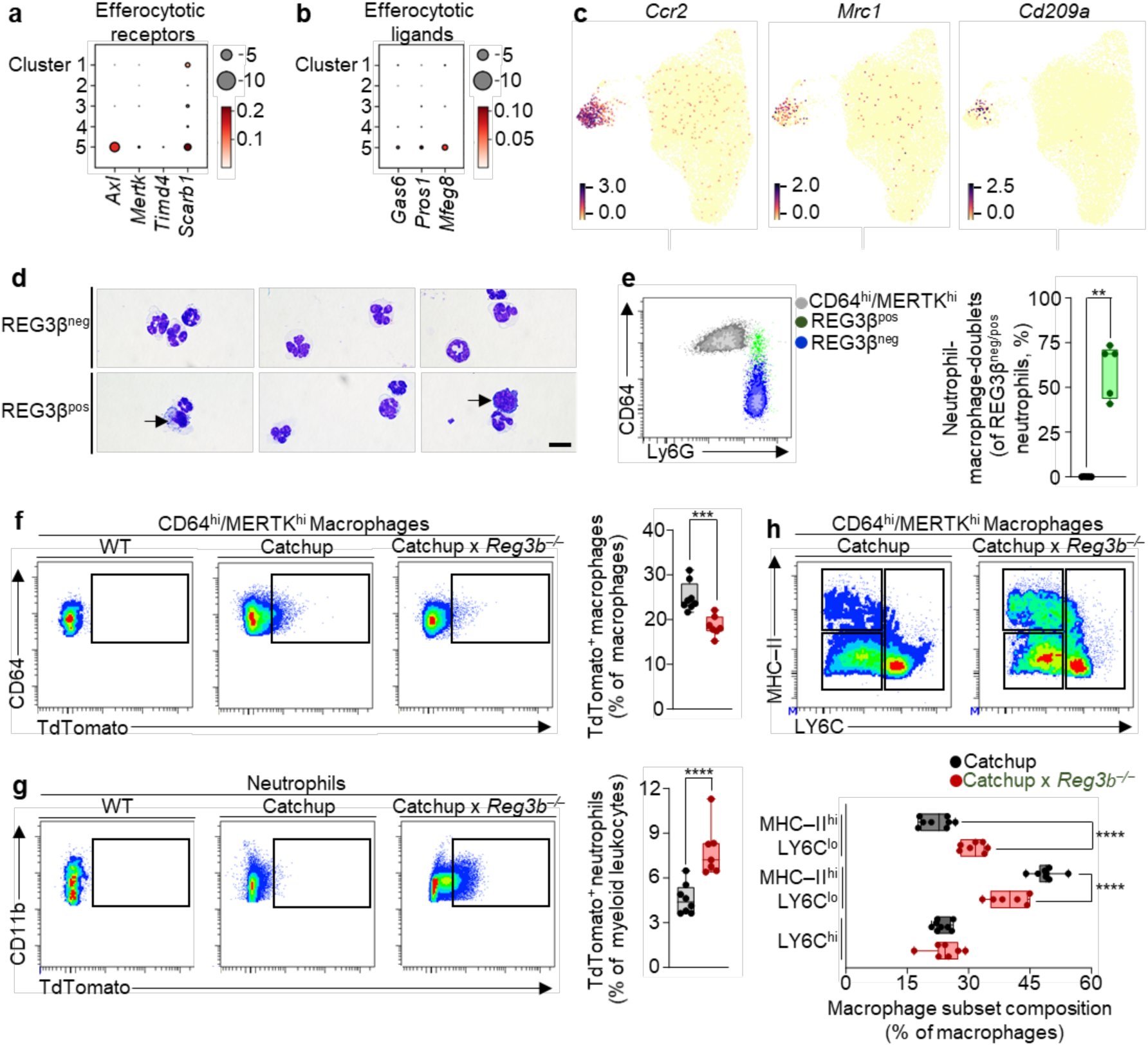
REG3β promotes neutrophil clearance via macrophage-mediated efferocytosis. **a**, **b**, Bubble plot showing gene expression of efferocytotic receptors and efferocytotic ligands across neutrophil clusters (see Fig.3). Bubble sizes represent percentage of cells expressing corresponding genes and bubble colors represents expression levels. **c**, Feature plot showing expression of *Ccr2*, *Mrc1*, and *Cd209a* across neutrophil clusters. **d**, Cytospin morphology of sorted REG3β^neg^ and REG3β^pos^ cardiac tissue neutrophils from WT mice 2 days after MI. Arrows indicate macrophages. Scale bars, 10μm **e**, Flow cytometry-based identification and quantification of macrophage-neutrophil-hybrid doublets obtained from WT hearts 2 days after MI. CD64^hi^/MERTK^hi^ macrophages (grey), REG3β^neg^ (blue) and REG3β^pos^ (green) neutrophils are shown. n = 5. **f**, Flow cytometric density plots and quantification of TdTomato^+^ CD64^hi^/MERTK^hi^ macrophages of Catchup (n = 8) and Catchup x *Reg3b^-/-^* (n = 7) hearts 2 days after MI. **g**, Flow cytometric density plots and quantification of TdTomato^+^ neutrophils of Ly6G^TdTomato^ (Catchup, n = 8) and Catchup x *Reg3b^-/-^* (n = 7) hearts 4 days after MI. **h**, Representative flow cytometric density plots and quantification of MHC-II^hi^/LY6C^lo^, MHC-II^lo^/LY6C^lo^, and LY6C^lo^ macrophage subset ratios in Catchup (n = 8) and Catchup x *Reg3b^-/-^* (n = 7) hearts 4 days after MI. Data are mean ± s.e.m.: Two-sided Mann Whitney test (**e**), two-sided unpaired t tests (**f**, **g**), one-way ANOVA followed by Sidak’s multiple comparison test (**h**). All experiments were conducted with male mice.

To functionally validate the role of REG3β for macrophage-mediated neutrophil clearance, we employed Ly6G^TdTomato^ mice, termed Catchup, which serve as reporter for neutrophils^41^. Analysis of Ly6G^TdTomato^ and Ly6G^TdTomato^//*Reg3b^-/-^*mice demonstrated a decreased ratio of TdTomato^+^ (CD64^hi^/MERTK^hi^) macrophages in infarct regions of Ly6G^TdTomato^//*Reg3b^-/-^*compared to Ly6G^TdTomato^ mice (Fig. 6f). Accordingly, ratios of non-phagocytosed TdTomato^+^ neutrophils were increased in Ly6G^TdTomato^//*Reg3b^-/-^*mice (Fig. 6g), indicating stimulation of efferocytosis by REG3β. Since efferocytosis promotes transition of macrophages to a more reparative phenotype, we analyzed the phenotype of macrophages based on differential surface expression of MHC-II and Ly6C in infarcted hearts of Catchup versus Catchup x *Reg3b^-/-^* mice^42,43,17^. We observed a substantial increase of proinflammatory MHC-II^hi^ /LY6C^lo^ and a decline of reparative MHC-II^lo^ /LY6C^lo^ macrophages in Ly6G^TdTomato^//*Reg3b^-/-^* compared to Ly6G^TdTomato^ mice. In contrast, the ratios of proangiogenic LY6C^hi^ macrophages were not affected (Fig. 6h). In conclusion, our findings demonstrate a crucial role of REG3β in orchestrating removal of neutrophil from infarcted hearts, thereby contributing to the termination of inflammation after MI and enabling proper cardiac remodeling.

## Discussion

Containment of inflammation after MI is critical for myocardial healing and proper remodeling, allowing damaged hearts to function despite the loss of contractile tissue. Neutrophils are important drivers of inflammation, arriving first at the scene after tissue damage, making they timely removal important to limit inflammation^44^. In addition, activated neutrophils exert direct adverse effects on the myocardium, potentially causing electrical storms, further emphasizing the need for their swift clearance^44,45^. We discovered that REG3β specifically removes aged and hyperactivated neutrophils, establishing a negative feedback loop within the heart to restrain tissue damaging-activities of neutrophils and terminate the first phase of inflammation. We postulate that this pathway establishes an active, tissue-instructed program of neutrophil elimination. So far, the function of cardiomyocytes in removal of neutrophils and limiting inflammation has not been recognized before. Attention was mostly focused on cardiac stromal cells and infiltrating macrophages, which also play important roles in restricting inflammation via different mechanisms^46,47^.

REG3β enriches at the interface of infarcted and non-infarcted regions of ischemically damaged hearts, where it presumably creates a barrier against further neutrophil expansion, protecting the viable myocardium. The scenario in infarcted hearts resembles the situation in the intestine, but for very different reasons. In the intestine, RegIIIγ acts as an antibacterial lectin to establish a zone that separates microbiota from the epithelial surface^48^. In both cases, REG proteins recognize the targets (hyperactivated neutrophils or microbiota) through the presence of mannose-containing glycoconjugates, arguing for an evolutionary conserved type of innate immune response^48,24^. Paucimannosylation has been broadly studied in lower organisms such as insects and nematodes but more recently also detected in vertebrates as an unconventional type of protein N-glycosylation^49^. Interestingly, human neutrophils contain large amounts of paucimannosidic glycans that are enriched in azurophilic granules but the role of paucimannosylation in disease processes has remained understudied^50^.

Specificity of REG3β for hyperactivated and aged neutrophils is achieved by enhanced translocation of paucimannosylated proteins from granules to the surface. Progressive loss of granule content is a hallmark of neutrophil activation and aging, although the functional relevance of the translocation is not fully understood^26,51^. We assume that the exposition of azurophil granules-derived paucimannosylated proteases, including MPO, and ELANE, on the surface of aged neutrophils serves a dual purpose: (i) facilitate migration of neutrophils through inflamed cardiac tissue and digestion of disposable extracellular material^52^; and (ii) label hyperactive neutrophils for removal by REG3β. After binding to paucimannosylated proteins REG3β enters neutrophils by clathrin-mediated endocytosis before accumulation within lysosomes and induction of lysosomal membrane permeabilization. We currently do not understand the exact molecular mechanism by which REG3β permeabilizes lysosomal membranes but it is tempting to speculate that a similar mode of action is employed as in bacteria. In bacteria, RegIIIα binds peptidoglycan carbohydrates on membranes before forming a hexameric membrane-permeabilizing oligomeric pore that kills the target^53^. Why REG3β does not form a pore within the cell membrane of neutrophils but presumably in the lysosomal membrane is unclear and requires further investigations.

REG3β initiates lysosome-mediated cell death for removal of activated and aged neutrophils, which differs from other ligand-dependent modes of killing, including pyroptosis, necroptosis, ferroptosis, NETOsis, and apoptosis^54^. We found that permeabilization of lysosomal membranes by REG3β releases cathepsins. Lysosomal membrane permeabilization and discharge of cathepsin are found in most types of cell death, including apoptosis, often amplifying ongoing cell death routines^55^. However, release of cathepsins does not necessarily lead to apoptosis but depends on the cellular context. We did not detect cleavage of the effector caspases-3/7, making it unlikely that activation of caspases is the principal executioner of cell death in REG3β-treated neutrophils. On the other hand, we observed higher ratios of the apoptotic marker AnnexinV on REG3β^pos^ neutrophils and enhanced display of phosphatidylserine, allowing efficient efferocytosis. The condition is reminiscent of a process coined lysoptosis, which occurs in the absence of caspase activation but requires cytoplasmic release of lysosomal cysteine peptidases^55^. Further studies are required to unravel the interactions between lysosomal cell death and apoptosis, although a mechanistic dissection is difficult given the rapid dynamics, complexity and crosstalk between individual cell death pathways^56^.

The discovery of REG3β-induced cell death of hyperactivated neutrophils paves the way for potential therapeutic interventions, allowing manipulation of neutrophil-driven inflammation, e.g. preventing neutrophil-dependent damage of cardiomyocyte membranes that cause arrhythmias and cell death^45^. Timing will be critical for such manipulations since hyperactivated neutrophils are not necessarily detrimental, acting in a highly stage-dependent manner. Exploitation of changes in glycoconjugates such as paucimannosylation may provide further tools for manipulation of specific subsets of neutrophils.

## Acknowledgements

We thank Dr. Ulrich Gärtner and Anika Seipp from Justus Liebig University Giessen for transmission and scanning electron microscopy. We thank Kikhi Khrievono for help with flow cytometry. The help of Kenny Mattonet in imaging experiments is greatly acknowledged.

