## Supplemental Figures and Methods for "Reg3β removes aged neutrophils after myocardial infarction"

### Analysis of human cardiac tissue

Human myocardial tissue specimens were obtained from the tissue bank of the Institute of Pathology, University Hospital Marburg. Inclusion criteria were a diagnosis of acute myocardial infarction within 48 hours prior to death. Autopsies were performed 24-72 hours post mortem according to standard institutional protocols. After macroscopic examination of the heart, the infarct territory was identified based on pallor, softening, hemorrhage, and loss of tissue elasticity. Tissue blocks were collected from (i) the infarct core, defined by confluent necrosis and myocyte dissolution, and (ii) the adjacent remote zone, defined as macroscopically intact myocardium located at least 1-2 cm outside the visibly infarcted region. Myocardial samples were trimmed to include full-thickness ventricular wall where feasible. Tissue was immediately immersion-fixed in 4% neutral-buffered formalin for ≥ 24 hours, routinely processed, and embedded in paraffin. Block identifiers, sampling location, and orientation were documented by the attending pathologist. Clinical characteristics were extracted from the electronic hospital information system. Use of human material was approved by the Ethics Committee of the Faculty of Medicine, Philipps University Marburg (reference 116/22). All procedures followed institutional guidelines, the Declaration of Helsinki, and applicable national regulations.

Anonymized tissue specimens were sectioned at 7 µm using a Thermo Scientific HM325 rotary microtome. Paraffin sections were deparaffinized in xylene (3 × 10 minutes), rehydrated through graded isopropanol (100%, 96%, 80%, 70%; 5–10 min each), and endogenous peroxidase was quenched in methanol containing 3% $H_2O_2$ for 30 minutes. Heat-induced antigen retrieval was performed in citrate buffer (pH 6.0) at 92–95 °C for 10 minutes. After rinsing, sections were circumscribed with a PAP pen (Z377821, Sigma Aldrich) and blocked in 5% BSA/PBS (30 min) followed by 1% BSA/PBS (5 min). Endogenous biotin was blocked using avidin-biotin reagents (20 minutes each; SP-2001, Vector Laboratories). Anti-REG3A (A5940, 1:300, Biotechne,) and anti-CD66b (305102, 1:400, Biolegend) were applied at working dilution and incubated overnight at 16°C The following day, sections were washed in distilled water and PBS and incubated with biotinylated secondary antibodies (anti-mouse IgM, 1:200, 715-065-140, Dianova and anti-mouse IgG, 1:200, 715-065-151, Dianova) for 45 minutes at 37 °C. After further washing and a brief block in 0.5% BSA/PBS, the avidin-biotin complex (PK-6100, Vector Laboratories) was applied for 30 minutes at 37°C. For detection of REG3A, tyramide (SAT700001EA, Perkin Elmer) signal amplification was performed by incubating sections with biotinylated tyramide for 20 minutes. Chromogenic development was carried out using 3,3′-diaminobenzidine (D12384, Sigma Aldrich) with ammonium nickel sulfate (09885, Sigma Aldrich). Prior to mounting, sections were counterstained with hematoxylin and eosin (H&E), dehydrated through graded ethanol (70%, 80%, 96%, 100%; 5 minutes each), cleared in xylene (3 × 10 minutes), and mounted with Eukitt (03989, Sigma Aldrich).

Immunostained sections were imaged using an Olympus AX70 microscope and a Zeiss Axio Imager.M2 microscope. Quantification of CD66b-positive cells was performed by an experienced examiner who manually counted all immunoreactive cells at 20× magnification using a systematic meandering scan across the full tissue section. Spatial association of CD66b-positive cells with REG3A expression was assessed on adjacent serial sections stained for REG3A.

**Animals**

Reg3b-deficient (*Reg3b*$^{-/-}$; B6;129-Reg2$^{tm1Lchr}$/H) mice were obtained from S. Hunt (University College of London). Neutrophil Ly6G-Cre; Rosa26tdTomato (Catchup; Ly6g$^{tm2621(cre)Arte}$) mice were bred with *Reg3b*$^{-/-}$ mice to generate *Catchup//Reg3b*$^{-/-}$ mice. Wild-type (WT) controls were C57BL/6J for studies using *Reg3b*$^{-/-}$ mice. All mice were kept in individually ventilated microisolator cages with 12-hour light-dark cycles in a specific pathogen-free facility with a temperature range between 20–24°C and humidity between 45-65%. Food (Altromin 1320, Altromin GmbH Lage), tap water, and nesting material were provided ad libitum. Experiments were performed in 8–14-weeks-old, age-matched male mice, since male C57/Bl6 mice show a higher incidence of heart failure and cardiac rupture after MI than female mice[57]. All animal experiments were performed in accordance with German animal protection laws and were approved by the local governmental animal protection committee, Regierungspräsidium Darmstadt, and followed relevant ethical regulations.

**Myocardial infarction**

Permanent LAD ligation was peformed as described with minor modifications[58]. Approximately 30 minutes before surgery, mice received buprenorphine (0.1 mg kg$^{-1}$, s.c.). Anaesthesia was induced with 3.0–4.0 vol% isoflurane in an optically shielded chamber, followed by orotracheal intubation and volume-controlled ventilation (tidal volume 1.5–1.9 ml; 60–80 breaths min$^{-1}$). Anaesthesia was maintained with 1.5–3.0 vol% isoflurane. Animals were positioned supine on a heated platform enabling continuous electrocardiogram monitoring, and ophthalmic ointment was applied. After confirming adequate anaesthetic depth, the left hemithorax was shaved and disinfected (povidone-iodine). An ~8 mm incision was made along the left midclavicular line, followed by blunt dissection to the chest wall. Bupivacaine (1 mg kg$^{-1}$) was administered locally before entering the third or fourth intercostal space. After blunt thoracotomy, the anterior cardiac surface was exposed, the LAD was identified, and, following a circumscribed pericardiotomy, either ligated in its mid-to-distal segment (distal to the diagonal branch) or at a proximal position (1–2 mm lower than the tip of the left auricle) for survival studies by using a 7-0 polyfil suture (K804H, Ethicon). The thoracotomy was closed in layers (intercostal, pectoral, skin). Isoflurane was discontinued at the end of the procedure, and mechanical ventilation continued until spontaneous respiration resumed after 3 to 5 minutes. Mice were then extubated and transferred to a heated recovery cage before being returned to their home cage once fully ambulatory. Postoperative analgesia consisted of buprenorphine (0.1 mg $kg^{-1}$, s.c.) at 4–6 h postoperatively and twice daily on postoperative days 1–2, supplemented with metamizole (200 mg $kg^{-1}$) administered via drinking water ad libitum. Animals were monitored closely and received additional buprenorphine as required.

**Cardiac MRI**

Cardiac MRI was performed on a 7.0 T Bruker Pharmascan system (Bruker BioSpin, Ettlingen, Germany) with 760 mT/m gradients, using a cryogenically cooled four-channel $^1$H phased-array receiver coil (CryoProbe) and a 72 mm volume resonator for transmission[58]. Cardiac self-gating and synchronization were achieved with IntraGate™. A gradient-echo sequence was applied for cine cardiac image acquisition (TR = 6.2 ms, TE = 1.3 ms, FOV = 2.2 × 2.2 $cm^2$, 1.0 mm slice thickness, 128 × 128 matrix, 20 cardiac phases). Two- and four-chamber long-axis views and 7-8 short-axis slices covering the left ventricle were collected. Mice were anesthetized with 1.5–2.0% inhaled isoflurane (1.0 L/min $O_2$/air) and maintained at 37°C using feedback-controlled warm-water circulation. Left ventricular volumes and function were quantified in Medis Suite QMass 3.2. (Medis Medical Imaging, Leiden, The Netherlands).

**Injections of Ly6G and REG3β antibodies**

Mice received intravenous tail vein injections 10 minutes before light sheet microscopy experiments (10µg of each antibody diluted in PBS in a total volume of 150µl per mouse). Neutrophil depletion was performed by intraperitoneal injection of Anti-Ly6G (clone: 1A8, Ab00295, Absolute Antibody, 50µg $d^{-1}$ diluted in 250µl PBS) on day 2, 3, 4, and 5 after infarct. Control mice were injected with IgG2a isotype control antibody (Ab00102, Absolute antibody, 50µg $d^{-1}$) at identical time points.

**Isolation and culture of neutrophils**

Bone marrow-derived neutrophils were collected from tibia and femur of mice. Bones were flushed with 5ml MACS buffer, composed of autoMACS rinsing solution (130-091-022, Miltenyi Biotec), supplemented with autoMACS BSA stock solution (130-091-376, Miltenyi Biotec), by using a 24 G needle and transferred to pre-chilled tubes. Peripheral blood neutrophils were obtained from whole blood drawn into EDTA tubes. Peritoneal neutrophils were obtained from the peritoneal cavity of mice after intraperiteonal injection of casein solution (9% w/v in PBS) for 2 consecutive days as described[59]. Thereafter, 5ml MACS buffer was intraperitoneally injected with a 24G needle, followed by withdrawal of the peritoneal fluid and transfer to pre-chilled tubes. Bone marrow, peripheral blood and peritoneal cell suspensions subsequently underwent red blood cell lysis by using ammonium chloride-based lysis buffer (46232S, Cell Signaling Technology), prior to purification by using the EasySep™ mouse neutrophils enrichment kit (#19762, StemCell). Cardiac cell suspensions from murine hearts were obtained by enzymatic digestion of excised hearts using Liberase Blendzyme (05401054001, Roche, 0.15 mg $ml^{-1}$) for 60 minutes at 37°C. Single cell suspensions were subsequently subjected to density gradient centrifugation using debris removal solution (130-109-398, Miltenyi Biotec), passed through a 30µm cell strainer, and resuspended in MACS buffer.

Primary murine neutrophils were cultured in Dulbecco's modified Eagle's medium (DMEM), high glucose, GlutaMAX™ Supplement, pyruvate (31966-021, Gibco) supplemented with 1% Pen/Strep at 37 °C with 5% CO2. 1.2 x $10^6$ murine neutrophils were aliquoted for stimulation experiments in 1 ml DMEM, high glucose, GlutaMAX™ Supplement, pyruvate, supplemented with 1% Pen/Strep at 37°C. Recombinant recombinant murine Granulocyte colony stimulating factor (G-CSF, 250-05, Thermofisher) at a concentration of 2ng $ml^{-1}$ and 100ng $ml^{-1}$ was additionally supplemented for short-term and for overnight stimulation experiments, respectively. Unless otherwise stated, stimulation experiments were carried out in 1.5ml culture tubes.

The following recombinant proteins were used in this study: recombinant murine REG3β (5110-RG, Biotechne), recombinant murine TNFα (AF-315-01A, Thermofisher), recombinant murine IFNγ (315-05, Thermofisher), recombinant PNGase F (P0709, New England Biolabs), recombinant Endo H (P0702, New England Bolabs), recombinant Endo F2 (P0777, New England Biolabs), and recombinant Endo D (P0742, New England Biolabs). LPS (00-4976-93) was purchased from Thermofisher. LLOme (L7393), D-Mannose (4440-M), and D-Galactose (D-3455) were obtained from Sigma Aldrich. All recombinant proteins, LPS, LLOme, and monosaccharides were resuspended and diluted in sterile PBS prior to stimulation experiments in vitro. The following pharmacological inhibitors were used in this study: Chlorpromazine hydrochloride (C8138, Sigma Aldrich), E-64d (E8640, Sigma Aldrich), Pepstatin A (J60237, Thermofisher), CA-074 Me (205531, Sigma Aldrich), and Nexinhib 20 (SML1919, Sigma Aldrich). All pharmacological inhibitors were diluted in DMSO (D5879, Sigma Aldrich) prior to adding to cultivated cells at indicated concentrations. Concentrations and duration of stimulation experiments with recombinant proteins, glycosidases, LPS, LLOme, monosaccharides, and pharmacological inhibitors are indicated for each experiment.

**Flow cytometry and cell sorting**

Single cell suspensions from infarcted areas and isolated neutrophils from bone marrow, peripheral blood and peritoneal cavity were stained in MACS buffer on ice for 30 minutes with the following antibodies: Anti-mouse CD45-FITC (clone: 30-F11, 0.25µg/test, 103108, Biolegend), anti-mouse CD11b-PE (clone: M1/70, 0.25µg/test, 101208, Biolegend), anti-mouse CD11b-FITC (clone: M1/70, 0.25µg/test, 101206, Biolegend), anti-mouse CD11b-PE (clone: M1/70, 0.25µg/test, 101208, Biolegend), anti-mouse Ly6G-FITC (clone: 1A8, 0.5µg/test, 127606, Biolegend), anti-mouse Ly6G-PE (clone: 1A8, 0.25µg/test, 127608, Biolegend), anti-mouse Ly6G-BV421 (clone: 1A8, 0.5µg/test 127628, Biolegend), anti-mouse CD64-APC (clone: X54-5/7.1, 0.5µg/test, 139306, Biolegend), anti-mouse CD64-BV421 (clone: X54-5/7.1, 0.5µg/test, 139309, Biolegend), anti-mouse CD54-BV421 (clone: YN1/1.7.4,

0.25µg/test, 116141, Biolegend), anti-mouse CD54-PE (clone: YN1/1.7.4, 0.1µg/test, 116107, Biolegend), anti-mouse TLR2-PE (clone: QA16A01, 1µg/test, 153003, Biolegend), anti-mouse CD63-APC (clone: NVG-2, 0.5µg/test, 143906, Biolegend), anti-mouse CD63-PE (clone: NVG-2, 0.5µg/test, 143903, Biolegend), anti-mouse TREM2-PE (clone: 6E9, 0.25µg/test, 824805, Biolegend), anti-mouse NOX2-PE (polyclonal, 0.25µg/test, BS-3889R, Thermofisher), anti-mouse CXCR4-BV421 (clone: L276F12, 0.25µg/test, 146511, Biolegend), anti-mouse CD62L-PE (clone: MEL-14, 0.25µg/test, 104408, Biolegend), anti-mouse CD49D-PE (clone: R1-2, 0.25µg/test, 103608, Biolegend), anti-mouse CD24-FITC (clone: M1/69, 0.25µg/test, 101805, Biolegend), anti-mouse Siglec-F-BV421 (clone: S17007L, 0.25µg/test, 155509, Biolegend), anti-mouse MERTK-PE (clone: 2B10C42, 0.5µg/test, 151506, Biolegend), anti-mouse MHC-II-APC-Cy7 (clone: M5/114.15.2, 0.25µg/test, 107628, Biolegend), anti-mouse Ly6C-PerCP-Cyanine5.5 (clone: HK1.4, 0.25µg/test, 128012, Biolegend), anti-human/mouse/rat MPO-FITC (polyclonal, MPO-112-FITC, 1µg/test, Thermofisher), anti-human/mouse/rat cardaic Troponin-AlexaFluor488 (clone: 990033, FAB18742G-100UG, 3µg/test, Biotechne), anti-mouse MMP-9-AlexaFluor488 (polyclonal, AF909G, 1µg/test, Biotechne), anti-mouse ADAM9-PE (polyclonal, AF949P, 1µg/test, Biotechne), and anti-mouse CD177-PE (clone: 1171A, FAB8186P-025, 2.5µg/test, Biotechne). Anti-mouse REG3β antibody (polyclonal, AF5110, 1µg/test, Biotechne) was directly labelled with APC for flow cytometric analysis using the Lightning-Link® APC Antibody Labeling Kit (705-0010, Biotechne). Anti-mouse ELANE (polyclonal, AF4517, 1µg/test Biotechne) and anti-mouse NGP (polyclonal, 600-401-GW9, 1µg/test, Thermofisher) were directly labelled with PE using Lightning-Link (R) R-PE Antibody Labeling Kit (703-0010, Biotechne). Neutrophils were defined as $CD45^{hi}$/$CD11b^{hi}$/$Ly6G^{hi}$ neutrophils and further divided into $REG3\beta^{neg}$ and $REG3\beta^{pos}$ neutrophil subsets. Macrophages were defined as $CD45^{hi}$/$CD11b^{hi}$/$Ly6G^{lo}$/$(CD64/MERTK)^{hi}$. Binding of externally added recombinant REG3β protein to cultured neutrophils was quantified by gMFI of REG3β on $REG3\beta^{pos}$ neutrophils via flow cytometry by using anti-mouse REG3β-APC conjugated antibody.

For intracellular expression analysis of REG3β, cardiac Troponin, MMP-9 and ADAM9 in cardiac tissue neutrophils, cells were fixated and permeabilized using the eBioscience Intracellular Fixation & Permeabilization Buffer Set (88-8824-00, Thermo Fisher) according to the manufacturer's protocol for staining of intracellular proteins. In order to compare viable and dead cell ratios of cell surface and intracellular $REG3\beta^{neg}$ and $REG3\beta^{pos}$ neutrophil subsets from cardiac tissue, Zombie Fixable Viability™ Sampler Kit (423117, Biolegend) was employed according to the manufacturer's instructions. For detection of paucimannosidic epitopes on cell surface of neutrophils, cells were first incubated with TruStain FcX™ PLUS antibody for 5 minutes on ice (clone: S17011E, 156603, 0.25µg/test, Biolegend) followed by incubation with AffiniPure® Fab Fragment Goat Anti-Mouse IgM, µ chain specific (polyclonal, 115-070-020,

1.5µg/test Jackson ImmunoResearch) on ice for 10 minutes to minimize non-specific binding. Thereafter, cells were incubated with Mannitou antibody (clone: Laz6-189, 0.5µg/test, Developmental Studies Hybridoma Bank) for 15 minutes on ice, followed by incubation with BV421-conjugated anti-mouse IgM antibody (clone: RMM1, 406517, 1µg/test, Biolegend) on ice for 15 minutes. Staining of the secondary antibody was used as control. Fluorescence compensation controls, fluorescence minus one controls (FMO), and isotype controls were used to assure correct compensation and gating. Flow cytometric analysis of leukocyte subsets was performed with the BD LSR fortessa cell analyser. Data were analyzed using the BD FACS Diva v6 Software. For calculation of cell numbers per mg of tissue, samples were normalized to the weight of tissues. The total number of neutrophils per ml blood was calculated by multiplication of total cells with the percentage of neutrophils. Sorting of $CD45^{hi}/CD11b^{hi}/Ly6G^{hi}/REG3\beta^{neg}$ and $CD45^{hi}/CD11b^{hi}/Ly6G^{hi}/REG3\beta^{pos}$ neutrophil subsets from infarcted hearts of WT was performed with the BD FACS Aria III cell sorter. Heart tissue preparation and staining was performed as described above.

**RNA sequence analysis and pathway annotation**

For RNA sequence analysis, RNA was isolated from sorted cardiac tissue $REG3\beta^{neg}$ and $REG3\beta^{pos}$ neutrophils by miRNeasy micro Kit (217684, Qiagen) combined with on-column DNase digestion (RNase-Free DNase Set, 79254, Qiagen) to avoid contamination by genomic DNA. RNA and library preparation integrity were verified with LabChip Gx Touch 24 (Perkin Elmer). RNA amounts were normalized and 10ng of total RNA was used as input for SMARTer® Stranded Total RNA-Seq Kit - Pico Input Mammalian (Takara Bio). Sequencing was performed on NextSeq2000 platform (Illumina) using P3 flowcell with 2 x 61bp paired-end setup. Trimmomatic version 0.39 was employed to trim reads after a quality drop below a mean of Q15 in a window of 5 nucleotides and keeping only filtered reads longer than 15 nucleotides[60]. Reads were aligned versus Ensembl mouse genome version mm39 (Ensembl release 109) with STAR 2.7.10a[61]. Alignments were filtered to remove: duplicates with Picard 3.0.0, multi-mapping, ribosomal, or mitochondrial reads. Gene counts were established with featureCounts 2.0.4 by aggregating reads overlapping exons on the correct strand excluding those overlapping multiple genes[62]. The raw count matrix was normalized with DESeq2 version 1.36.0[63]. Contrasts were created with DESeq2 based on the raw count matrix. Genes were classified as significantly differentially expressed at average count > 5, multiple testing adjusted p-value < 0.05, and -0.585 < log2FC > 0.585. The Ensemble annotation was enriched with UniProt data (Activities at the Universal Protein Resource (UniProt)). All downstream analyses are based on the normalized gene count matrix. Dimension reduction analyses (PCA) were performed on regularized log transformed counts using the R packages FactoMineR[64]. DEGs were submitted to gene set overrepresentation analyses with KOBAS [65]. The resulting bubble plot shows pathways with Benjamini-Hochberg corrected p-value < 0.05 (represented by dashed line). The larger gray circles are scaled to the number of genes comprising the respective pathway, while the smaller colored circles represent subsets found to be DEGs.

Transcriptome data of cardiac tissue REG3β$^{neg}$ and REG3β$^{pos}$ neutrophils have been deposited at NCBI Gene expression omnibus (GSE312422, https://www.ncbi.nlm.nih.gov/geo/query/acc.cgi?acc=GSE312422).

**Single cell RNA sequence analysis of cardiac neutrophils**

Single-cell suspensions were quantified and processed using the Rhapsody Scanner (BD Biosciences) in accordance with the manufacturer's standard protocols for cell loading and library construction. Sequencing was performed on the Illumina NextSeq 2000 platform. Raw sequencing reads were aligned to the Mus musculus reference genome (mm39) and preprocessed with the BD Biosciences cwl-runner pipeline, generating data in the annotated data format for downstream analysis.

Subsequent computational analyses were conducted using Scanpy[66]. Quality control procedures considered both the number of genes detected per cell and the proportion of mitochondrial transcripts. Cells expressing fewer than 500 genes or exhibiting mitochondrial transcript content above 37% were excluded (n = 1,866). Additionally, 12,027 genes detected in fewer than 30 cells (< 0.01%) were filtered out. Expression data were scaled relative to the median library size of all cells and transformed to logarithmic space to stabilize dispersion. Principal component analysis (PCA) was then applied, and the leading 50 components were selected for downstream computations. Further analyses, including Uniform Manifold Approximation and Projection (UMAP) embedding and community-based clustering, were performed on the PCA-reduced space [67,68]. Final visualization and interactive data exploration were carried out using the CellxGene platform (https://cellxgene.cziscience.com). Density plots of REG3β$^{neg}$, Pan and REG3β$^{pos}$ cardiac tissue neutrophils were calculated by gaussian kernel density estimation implemented in Scanpy's "embedding_density" function[66]. We utilized the sc-framework toolkit for data processing and pseudotime calculation, which integrates the PALANTIR package [69,70]. This algorithm models the trajectories of differentiating cells by treating their transcriptomes as probabilistic processes and using entropy to measure their plasticity along the trajectory. Terminal states were predicted, cells were assigned to branches and ordered along a high-resolution pseudo-time. Scores of biological processes were calculated by summing up all counts for defined gene sets for each cluster. Corresponding gene signatures for calculating bone marrow proximity, aging, inflammatory response, metallopeptidase activity, ROS production, and phagocytosis scores are provided in (Suppl. Table 1).

Single cell transcriptome data of cardiac tissue REG3β$^{neg}$ and REG3β$^{pos}$ neutrophils are available online at NCBI Gene expression omnibus (GSE312422, https://www.ncbi.nlm.nih.gov/geo/query/acc.cgi?acc=GSE312422).

**N-glycan analysis of the cell surface of neutrophils**

For isolation of membrane proteins, sorted neutrophils were lysed in Triton lysis buffer (10 mM Tris-HCl, 150 mM NaCl, 1 mM EDTA, 1% (*v*/*v*) Triton X-114) containing protease inhibitor cocktail. Cell suspension was sonicated and incubated at 4°C overnight. Following centrifugation, the supernatant was overlayed onto sucrose cushion (6% (*w/v*) sucrose, 10 mM Tris-HCl, 150 mM NaCl, 0.06% Triton X-114) and incubated at 37°C for 20 minutes leading to clouding of the solution (micellization). Micelles were separated into two phases by centrifugation. The upper aqueous phase was removed and micellization was repeated twice with the remaining lower detergent phase and kept on ice. Subsequent protein precipitation of the detergent phase containing membrane proteins was performed by gradually adding four volumes of methanol, two volumes of chloroform and three volumes of ultra-pure water. Upon centrifugation, liquid interphase containing proteins were again precipitated by adding three volumes of methanol, centrifuged and samples were dried in a vacuum concentrator.

Dried protein samples were denatured via resuspension in 1.33% SDS (*w/v)* and incubated for 10 minutes at 65°C. Subsequently, 4% (*v*/*v*) Igepal-CA630 was added and gently shaken for 15 minutes at room temperature. Deglycosylation was performed by adding 1.2 U of PNGaseF in 5 × PBS and overnight incubation at 37°C. Released N-glycans were labeled with procainamide hydrochloride (ProA, . ProA (172.8 mg $mL^{-1}$) dissolved in 30% (*v*/*v*) glacial acetic acid in DMSO) for 1 hour at 65°C. The reducing agent, 2-picoline borane (179.2 mg mL-1, dissolved in 30% (*v*/*v*) glacial acetic acid in DMSO) was subsequently added and incubated for 1.5 hours at 65°C. To remove the free label and reducing agent, hydrophilic interaction liquid chromatography - solid phase extraction (HILIC-SPE) was used. Samples were next added to the wells and washed with 96% (*v*/*v*) acetonitrile (CAN). ProA labeled and cleaned-up N-glycans were eluted twice with ultra-pure water, and both eluates were combined. Enrichment of labeled N-glycan was performed using cotton ziptips. Samples were applied to columns and centrifugated. The samples were then eluted with ACN containing 100 mM ammonium formate. Following centrifugation, samples were transferred and analyzed by HILIC on an Acquity H-class ultra-high-performance liquid chromatography (UHPLC) system (Waters) consisting of a quaternary solvent manager, sample manager, column oven, and a fluorescence detector (FLD) set at excitation and emission wavelengths to 310 and 370 nm, respectively. ProA labeled N-glycans were separated on a Waters bridged ethylene hybrid (BEH) glycan chromatography column, 150 × 2.1 mm, 1.7 µm, BEH particles, with 100 mmol $L^{-1}$ ammonium formate, pH 4.4, as solvent A and ACN as solvent B. Separation was achieved by a linear gradient of 70–53% ACN at a flow rate of 0.56 mL $min^{-1}$ in a 25 minutes analytical run. Obtained chromatograms were separated into 28 chromatographic peaks and their abundance was expressed as percentage of total integrated area. N-glycans were annotated according to measured *m/z* value and recorded fragmentation spectra obtained by Bruker

Compact Q-ToF mass spectrometer coupled to UHPLC via Ion Booster ion source and controlled using HyStar software (Bruker Daltonics). All recorded spectra were analyzed using Bruker Data Analysis software version 4.4 (Bruker). N-glycan compositions and structural features were determined using GlycoMod and GlycoWorkbench software tools[71].

**Glycoproteome analysis of neutrophils**

Proteins from neutrophil pellets were extracted with 4% SDS, 100mM Tris, pH 7.6 and protease inhibitors (04693116001, Roche). Protein purification was performed by 80% acetone precipitation. Proteins were re-solubilized in 6M urea/2M thiourea (after reduction with 10mM dithiothreitol (DDT) and alkylation with 55mM odoacetamide (IAA)), digested with Lysyl endopeptiase (LycC, 125-05061, Fujifilm) (ratio 1:10) at room temperature for 3 hours, and then digested with trypsin (37283.03, Serva) (ratio 1:50) at 37°C overnight. A C18 stage was used for peptide desalting prior to liquid chromatography-tandem mass spectrometry (LC-MS/MS) analysis. In-house made (Hydrophobic Interaction Liquid Chromatography) HILIC and Concanavalin A (ConA) tips were used to enrich glycopeptides. Peptides were separated on a C18 capillary column (150 mm x 1.7 µm x 75 µm, ReproSil-Pur 120 C18-AQ) using a 150-minute gradient of acetonitrile with 0.1% formic acid. Orbitrap Q-exactive HF was used for mass spectrometry analysis. Samples were measured using the Data dependent workflow analyzing major 15 ions (TOP 15 DDA workflow) in positive Electrospray ionization (ESI) mode. A resolution of 60,000 was used for MS measurements and 15,000 MS/MS measurements. Ions with charge state +1-+7 in range of m/z: 350-1800 were used for MS/MS fragmentation using a normalized collision energy (NCE) of 32. Peptides with charge states of +1, +2, and >+8 were excluded from fragmentation and dynamic exclusion was set to 30 seconds. The collected data were processed using Byonic software (Protein Metrics). Neutrophils protein digests for proteomic analysis were prepared as described above. Mass spectrometry methodology used for proteomic analysis was slightly modified as follows: Ions with charge state +2-+7 in range of m/z: 350-1800 were used for MS/MS fragmentation using an NCE of 28. Further analysis of the collected data was carried out using the Maxquant software tool.

**Neutrophil viability and cytotoxicity assays**

Viability, death and phosphatidylserine externalization of neutrophils were monitored by using the Annexin V apoptosis detection kit with 7-AAD (640922, Biolegend), according to the manufacturer's instructions and analyzed via flow cytometry. Detection of neutrophil cell death in real time was performed by using the IncuCyte®. For application of the Cytotox Green Assay, neutrophils were seeded at a cell density of 250,000 $ml^{-1}$ in fibronectin (341631, Sigma Aldrich) pre-coated 24 well plates and cultured in the IncuCyte® S3 Live-Cell Analysis System (Sartorius). Cytotox green reagent (4633, Sartorius, 250nM) and recombinant proteins were simultaneously added. Four phase contrast and green fluorescent images per well were recorded every hour throughout the assay. Live-cell images were subsequently processed and analyzed for number of green-positive cells by using IncuCyte® integrated analysis software (IncuCyte ZOOM version 2016A, Sartorius). Monitoring of cell death was further determined by measuring the loss of fluorescence of neutrophils preloaded with MitoSpy™ Red CMXRos (424801, Biolegend). Proportions of $MitoSpy^{low}$ neutrophils were identified and quantified by flow cytometric analysis. The release of lactate dehydrogenase as a measure of cytotoxicity was determined in supernatants of neutrophil cell cultures using the CytoTox 96 Non-Radioactive Cytotoxicity Assay (G1780, Promega) according to the manufacturer's instructions. Intracellular ROS production was detected by using the cell permeable dye Dihydrorhodamine 123 (DHR 123, D23806, Thermofisher). Neutrophils were loaded with 150µM DHR123 for 15 minutes prior to stimulation experiments and oxidation of DHR123 was measured and quantified as gMFI of DHR123 via flow cytometry.

**Analysis of neutrophil phagocytotic activity and neutrophil activation**

Phagocytosis of cardiac tissue neutrophils was analyzed by using pHrodo™ Phagocytosis Particle Labeling kit (A10026, Thermofisher) according to the manufacturer's instructions and quantified as gMFI of pHrodo via flow cytometry. Phagocytotic uptake of cardiomyocyte-derived cardiac Troponin in cardiac tissue neutrophils was determined and quantified as gMFI of cardiac Troponin by using anti-human/mouse/rat cardiac Troponin-AlexaFluor488 antibody in flow cytometric analysis. Bone marrow derived murine neutrophils (BMN) were stimulated with (i) LPS (200ng $ml^{-1}$) for 2 hours to obtain activated neutrophils, (ii) with PBS for 24 hours to acquire an in vitro aged phenotype, and (iii) with LPS (200ng $ml^{-1}$) for 24 hours to generate aged neutrophils with an activated phenotype. BMN treated with PBS for 2 hours were defined as basal neutrophils. Flow cytometry was employed to determine activation and aging of neutrophils by using dihydrorhodamine 123 (DHR123), CD54, CXCR4, and CD62L.

**Caspase 3/7 activity and neutrophil lysosome membrane permeabilization assays**

Caspase 3/7 activation in neutrophils was performed by using the CellEvent Caspase-3/7 Green Flow Cytometry Assay Kit (C10427, Thermofisher) according to the manufacturer's protocol. Lysosomes of neutrophils were initially labelled with Lysotracker probes (50nM, L12492, Thermofisher) according to the manufacturer's protocol. Lysosomal membrane permeabilization of cultured neutrophils was subsequently quantified as the percentage of Lysotracker-negative neutrophils via flow cytometry.

**Light sheet fluorescence microscopy**

Three-dimensional imaging of infarcted hearts via light sheet microscopy was adapted from the BALANCE protocol[72]. The following antibodies were used: Anti-mouse Ly6G-AF647 (clone: 1A8, 127610, Biolegend), and anti-mouse REG3β (polyclonal, AF5110, Biotechne). Anti-mouse REG3β was directly labelled with AF790 using Alexa Fluor™ 790 Antibody Labeling Kit (A88070, Thermofisher). Ten minutes after intravenous injection of antibodies, mice were sacrificed and perfused with PBS. Hearts were excised and fixated in 4% paraformaldehyde

(PFA) for 2 hours at room temperature. Afterwards, hearts were dehydrated in an ascending ethanol series in ddH2O (v/v) of 30%, 50%, 70%, 90% and two times 100% (9065.2, Roth) for at least 4 hours at 4°C under agitation. Hearts were next bleached in 5% (v/v) hydrogen peroxide and 5% (v/v) DMSO in 100% ethanol for 4 hours at 4°C. After washing in 100% ethanol for 1 hour, samples were warmed up to room temperature and transferred in pure ethylcinnamate (ECi, 112372, Sigma Aldrich) for at least 4 hours prior to imaging. Light sheet imaging was performed with the Ultramicroscope II and ImSpector software (both LaVision BioTec). 3D reconstruction was done using Imaris Software (Bitplane).

**Histology and immunofluorescence**

Hearts were fixed with 3% PFA for 2 hours at room temperature. After fixation, hearts were transferred to 15% sucrose solution for 2 hours at room temperature, followed by overnight incubation in 30% sucrose (9097.1, Roth) at 4 °C, and then embedded in Tissue-Tek® O.C.T (SA1467N, Scienceservices). Cryosections were 10 µm thick, and 6–8 regions were sampled from each heart. Histological examination of heart tissue was performed after hematoxylin and eosin staining (H&E, Chroma/Waldeck), following standard protocols[73]. Slices were thawed for 30 minutes at room temperature and washed with PBS. Prior incubation with the primary antibodies anti-mouse MMP9 (AF909, Biotechne), anti-mouse ELANE (AF4517, Biotechne), and anti-mouse MPO (AF3667, Biotechne), antigen retrieval was performed as follows. Sections were incubated in 10mM citrate buffer (pH 6) for 20 minutes at 95 to 99°C. After cooling down, slices were permeabilized for 10 minutes with 0.1% Triton X-100, and blocked with 3 % BSA solution for 1 hour at room temperature. Primary antibody incubation was performed overnight at 4 °C with the following antibodies, diluted in PBS containing 1% BSA: Anti-mouse Ly6G-AF647 (1:100, clone: 1A8, 127610, Biolegend), anti-mouse MMP-9 (1:200, AF909, Biotechne), anti-mouse ELANE (1:200, AF4517, Biotechne), and anti-mouse MPO (1:200, AF3667, Biotechne). Infarcted cardiac tissue slices were additionally incubated with Phalloidin-FITC (1/100, F432, ThermoFischer) to distinguish infarcted from non-infarcted tissue. After three PBS washes, Alexa-conjugated secondary antibodies were applied for 1 hour at room temperature including chicken Anti-goat AF594 (1:200, A21468, Invitrogen) and donkey Anti-sheep AF594 (1:200, A11016, Invitrogen). Nuclei were counterstained using 4′,6-Diamidin-2-phenylindol (DAPI, 1:2000, D9542, Sigma Aldrich) for 10 minutes at room temperature. Stained slides were finally washed with PBS and coverslips were mounted using Mowiol mounting medium (composed of 2.4g Mowiol 4-88, 0713.1, Roth; 6g Glycerol, 3783, Roth; 6ml ddH2O; 12ml 0.2M Tris-HCl, pH 8.5, 9090, Roth). Fluorescent images were acquired on a Leica TCS SP8 confocal microscope, and brightfield images on a Leica Thunder Imager microscope, both controlled with LAS X Life Science software.

Peritoneal neutrophils were attached to slides using a cytospin 4 centrifuge (ASHA78300003, ThermoFischer) at 50.000 cells per slide. Slides were immediately fixed with 4% PFA for 15 minutes at room temperature. After washing, cells were blocked with 5% serum (matching antibody host species) and 1% BSA for 1 hour at room temperature. Neutrophils were then stained overnight at 4 °C with antibodies diluted in 0.5 % serum plus 0.5 % BSA. The following antibodies were used: Anti-mouse MPO (1:200, AF3667, Biotechne), anti-mouse ELANE (1:200, PA5-88916, Invitrogen), Anti-mouse REG3β (1:100, clone: 518630, MAB5110, Biotechne), and anti-mouse NGP (1:300, 600-401-GW9, Thermofisher). Cells were additionally incubated with WGA-FITC (1:100, L4895, Merck) to visualize the plasma membrane. After three PBS washes, cell surface paucimannose staining was performed by incubating cells with undiluted Mannitou antibody supernatant (clone: Laz6-189, Developmental Studies Hybridoma Bank) for 30 minutes at room temperature. Afterwards, cells were incubated with the appropriate secondary antibodies: chicken anti-rat-AF647 (1:200, A21472, Invitrogen), chicken anti-goat-AF594 (1:200, A21468, Invitrogen), goat anti-rabbit AF594 (1:200, A11012, Invitrogen) and goat anti-mouse IgM (1:200, A21044, Invitrogen). After washing, samples were incubated with DAPI (1:2000) and mounted as described above. Specificity of antibody staining was assessed by incubating slides with secondary antibodies alone to distinguish true primary antibody signal from background binding. Imaging was performed on a Zeiss 880 Airyscan confocal microscope using a LD LCI Plan-Apochromat 63x/1,2 Imm Korr DIC M27 objective, and is equipped with a 32-channel Airyscan detector, enabling image processing with Airyscan functionality in Zen Black (Zeiss).

**Expansion microscopy**

Peritoneal neutrophils were attached to 10 mm round coverslips (0112500, Marienfeld Superior) as described above. To analyze protein distribution at super-resolution in cells, an immunostaining and expansion microscopy protocol was adapted[74]. Immunostaining was performed as described above except for 2 modifications, (i) permeabilization was performed with PBS containing 0,4% Triton X-100 for 5 minutes, and (ii) incubation times for antibodies were extended for up to 24 hours for primary antibodies and overnight for secondary antibodies, both at 4 °C. The following antibodies were used: Anti-mouse LAMP1 (1:100, clone 747203, MAB4320, Biotechne), anti-mouse REG3β (1:50, AF5110, Biotechne), anti-Galectin-1 (1:100, AF1245, Biotechne), anti-mouse Cathepsin D (1:200, AF1029, Biotechne) and anti-mouse Cathepsin B (1:100, AF965, Biotechne). Secondary antibodies we used: donkey anti-rat AF594 (1:100, A21209, Invitrogen), rabbit anti-goat ATTO 647N (1:100, 605-456-013S, Rockland), and rabbit anti-sheep ATTO 647N (1:100, 3321, Hypermol). Imaging was performed on a Zeiss 880 Airyscan confocal microscope using an LD LCI Plan-Apochromat 63x/1,2 Imm Korr DIC M27 objective.

**Image processing and analysis**

Immunofluorescent images were processed in ImageJ, v1.54r. Contrast and brightness were adjusted using the 5D contrast optimizer plugin, applied in the same way to all images with the same staining to avoid bias. To quantify neutrophils in the infarct and border zones, we first outlined the infarct zone manually using the Phalloidin channel in ImageJ and defined as region of interest (ROI). In parallel, neutrophils were identified from the Ly6G channel using a Cellpose model, which generated a mask marking all neutrophils. We wrote an ImageJ macro to combine the ROI with the Cellpose mask, allowing automated counting of neutrophils inside the infarct zone. For non-infarcted remote zone, the macro measured neutrophil counts in 100 µm steps moving outward from the infarct edge, covering a total distance of 1000 µm. We also counted the total number of neutrophils in each section as a reference. All measurements were exported to an Excel sheet using a small Python macro to organize the data. For co-localization analysis between EEA1 and REG3β, as well as LAMP1 and REG3β, we used the JacoP plugin in Image J and quantified co-localization using the Pearson coefficient.

**Transmission and scanning electron microscopy**

Neutrophils were fixated with 4% PFA and incubated overnight at 4°C. Post-fixation was performed in 1 % aqueous osmium tetroxide followed by dehydration through series of graded alcohol (25%, 50%, 70%, 90% and 100%). Afterwards, samples were embedded by using the Agar 100 Resin kit (AGR1031, Agar Scientific), cut in ultrathin sections via ultramicrotomy (Ultramicrotome, Leica Microsystems) and imaged by using a transmission electron microscope (Zeiss EM902). Neutrophils were fixated in 2.5 % glutaraldehyde and incubated overnight at 4°C. Post-fixation was performed in 1 % aqueous osmium tetroxide followed by dehydration through series of graded alcohol (25%, 50%, 70%, 90% and 100%). Afterwards, samples were dried by critical point $CO_2$ treatment, gold sputter-coated and examined using the Phillips XL30 SEM (Philips Electron Optics) [75].

**Protein extraction and immunoblot analysis**

Proteins from tissues and neutrophil cultures were extracted by using lysis buffer (0.1 M Tris-HCl pH 8.8, 0.01 M EDTA, 0.04 M DTT, 10 % SDS, pH 8.0) supplemented with protease inhibitors (500 $\mu$g ml$^{-1}$ Benzamidin, 2 $\mu$g ml$^{-1}$ Aprotinin, 2 $\mu$g ml$^{-1}$ Leupeptin, 2 mM PMSF, 1 mM Sodium Vanadate, 20 mM Sodium Fluoride). Separation of proteins were performed by SDS-PAGE on Gradient NuPAGE 4–12% Bis-Tris gels (NP0321BOX, Thermofisher) followed by blotting onto nitrocellulose membranes (88018, Thermofisher). Membranes were probed with the following specific primary antibodies: Anti-mouse REG3β antibody (polyclonal, AF5110, 1:500, Biotechne), anti-mouse NGP (polyclonal, 600-401-GW9, 1:2000, Thermofisher), anti-human/mouse MPO (polyclonal, AF3667, 1:2000, Biotechne), anti-mouse ELANE (polyclonal, AF4517, 1:2000, Biotechne), anti-mouse MMP-9 (polyclonal, AF909, 1:1000, Biotechne), anti-mouse ITGAM (clone: 908649, MAB11241, 1:2000, Biotechne) and anti-mouse CD177 (clone: E1V7N, 28740, 1:2000, Cell Signaling Technology). Secondary antibodies including anti-sheep IgG horsereadish peroxidase-conjugated polyclonal antibody (HAF016), anti-rabbit IgG horsereadish peroxidase-conjugated polyclonal antibody (catalogue no. HAF008), anti-goat IgG horsereadish peroxidase-conjugated polyclonal antibody (catalogue no. HAF017), and anti-rat IgG horsereadish peroxidase-conjugated polyclonal antibody (catalogue no. HAF005) were obtained from Biotechne. Immunoreactive proteins were visualized by chemiluminescence using SuperSignal™ West Femto Maximum Sensitivity Substrate (34095, Thermofisher) and a ChemiDoc™ MP Imaging System (Bio-Rad). Image Lab software 5.0 (Bio-Rad) was employed for densitometric quantification for band intensities.

**Statistical analysis**

Data were analyzed using GraphPad PRISM 10. Normality for group samples with $n \geq 6$ was assessed by using the Shapiro-Wilk normality test. Data obtained from group sizes ≥ 6 that passed the normality test were analyzed by parametric tests including two-sided unpaired t tests for two groups, one-way ANOVA followed by Sidak's multiple comparison test for more than two groups, and two-way ANOVA followed by Tukey's multiple comparison or Sidak's multiple comparison test for comparison of multiple conditions to two or more groups. Group sizes of $n<6$ were analyzed using non-parametric tests, including nonparametric Kolmogorov-Smirnov test and Two-sided Mann Whitney test for two groups, and Kruskal-Wallis 1-way ANOVA followed by Dunn's multiple comparison test for more than two groups. Kaplan-Meier survival curves were compared by the log-rank (Mantel-Cox) test. Correlation analysis was performed by using Pearson`s test. Statistical significance was defined as $P < 0.05$, with significance levels represented as $^{*}P < 0.05$, $^{**}P < 0.01$, $^{***}P < 0.001$ and $^{****}P < 0.0001$. Power calculations were not applied to predetermine sample sizes but are equivalent to group sizes reported in literature. All obtained data were included in our analyses except for animal health concerns or in case of statistically based outlier identification.

### SUPPLEMENTAL FIGURES

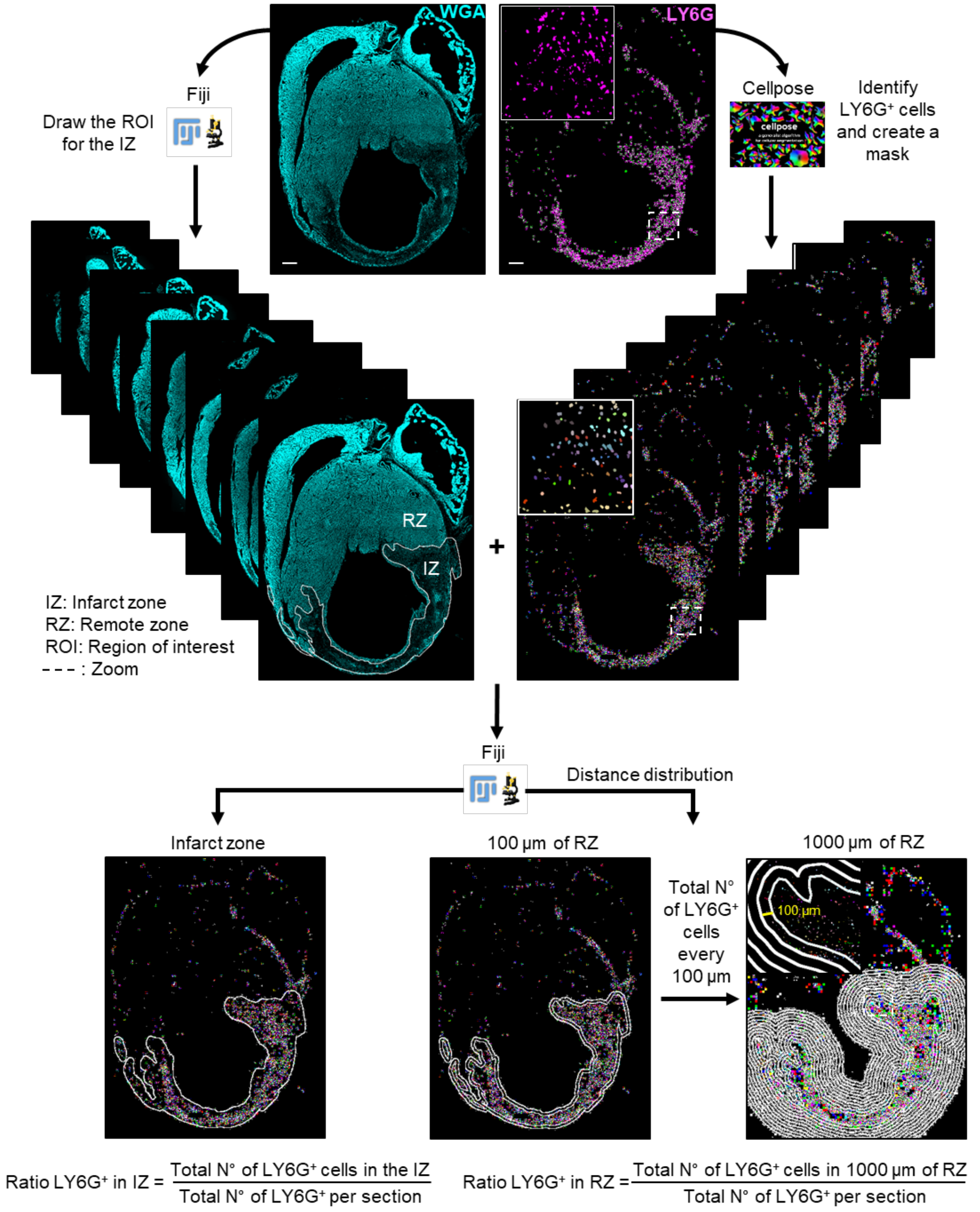


**Supplemental Figure 1: Workflow for identification and quantification of neutrophils within infarcted and non-infarcted remote heart regions.**

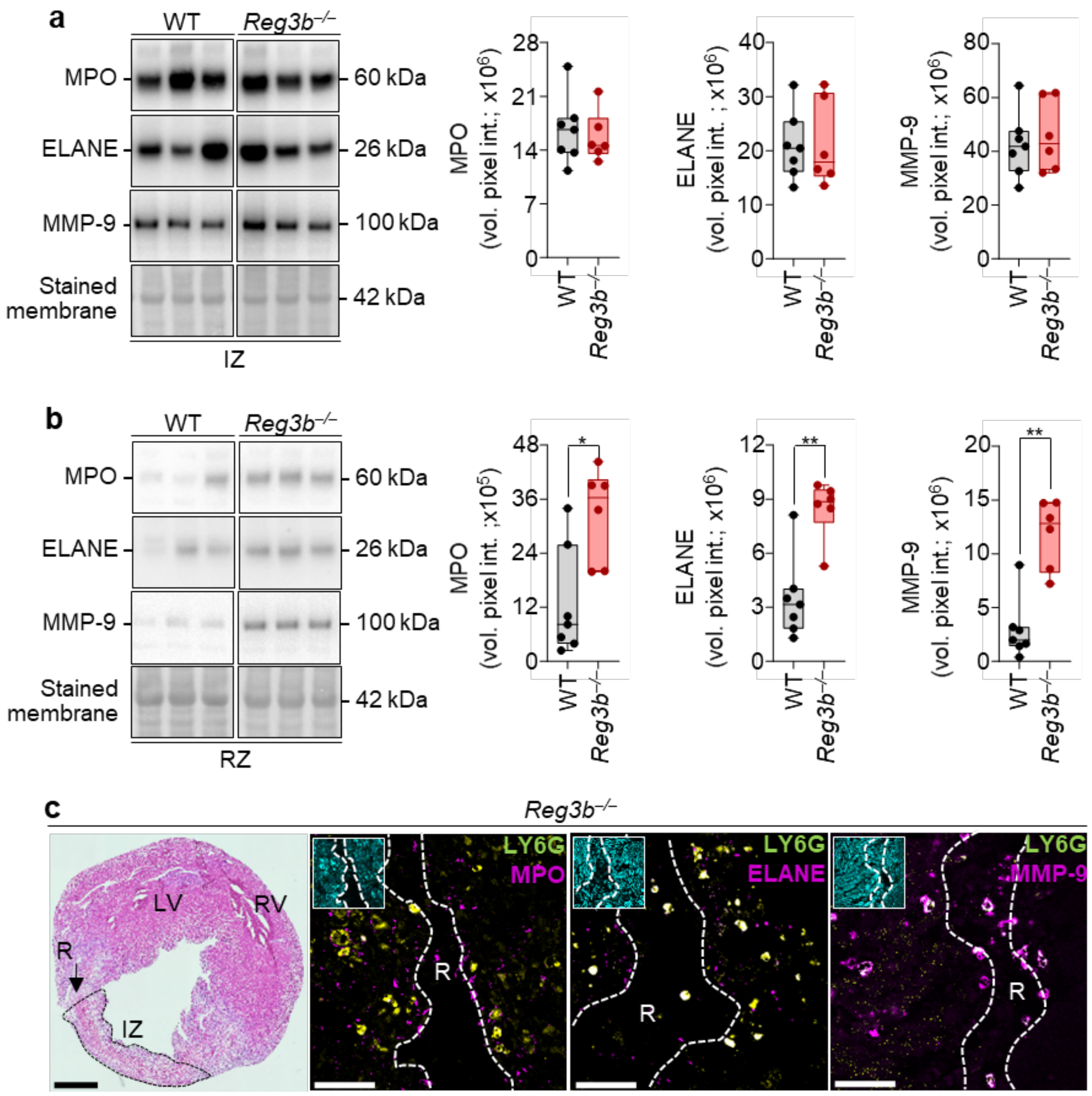


**Supplemental Figure 2: Distribution of neutrophil-derived proteases in infarcted WT and Reg3b-deficient hearts. a**, **b**, Immunoblots and semi-quantitative analysis based on the mean volume pixel intensity (vol. pixel int.) of Myeloperoxidase (MPO), Neutrophil elastase (ELANE), and Matrix metalloproteinase-9 (MMP-9) within infarcted zone (IZ) and non-infarcted remote zone (RZ) of wild-type (WT) and *Reg3b* deficient (*Reg3b*$^{-/-}$) mice 4 days after infarct. WT, n = 7 and *Reg3b*$^{-/-}$, n = 6. RedAlert staining demonstrates equal sample loading. **c**, Hematoxylin and eosin (HE) and immunofluorescence (IF) staining of LY6G$^{+}$ neutrophils (green), MPO, ELANE, and MMP-9 (all in magenta) of a ruptured *Reg3b*$^{-/-}$ heart. Left ventricle (LV), right ventricle (RV), IZ, and position of cardiac rupture (R) is indicated. Scale bars, 100µm (HE) and 10µm (IF). Data are mean ± s.e.m. Two-sided Mann Whitney test. All mouse experiments were conducted with male mice.

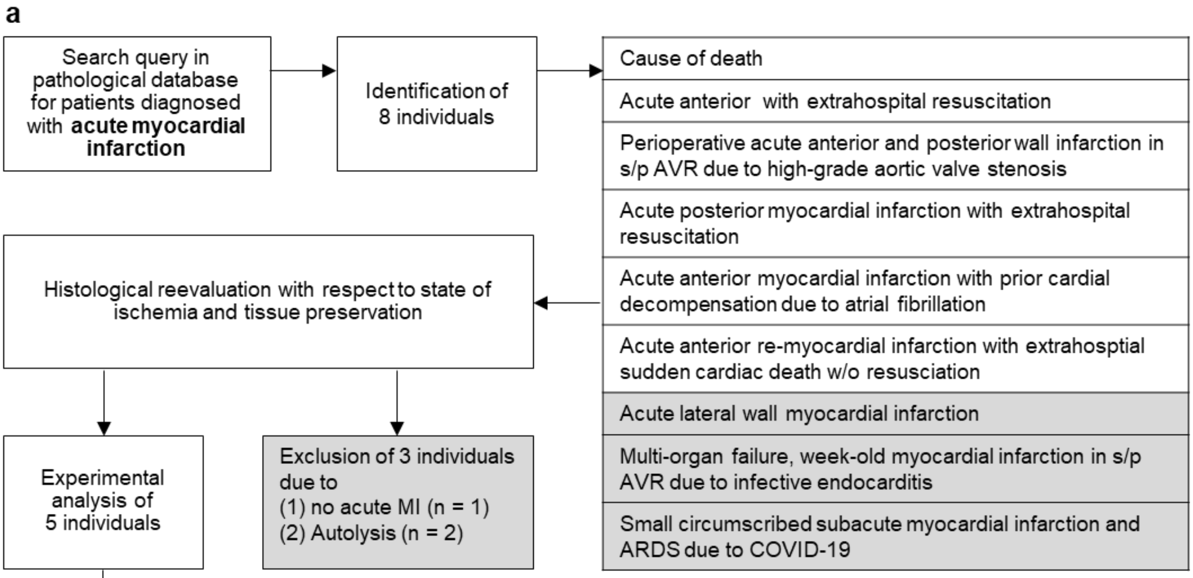


| Characteristics | |
|---|---|
| Median Age, years | 63 |
| Sex<br>Male<br>Female | <br>5<br>0 |
| Median time from death to autopsy, days | 2 |
| Mean Body Mass Index, kg $m^{-2}$ | 29.4 |
| Arterial hypertension, n | 4 |
| Diabetes mellitus, n | 2 |
| History of smoking, n | 2 |
| Dyslipidemia, n | 3 |
| Coronary artery disease, n | 3 |
| Prior Resuscitation, n | 2 |
| Prior Intensive Medical Care, n | 3 |
| Prior extracorporeal life support use, n | 2 |
| Prior Cardiac Surgery, n | 1 |
| Mean heart weight, g | 518 |
| Left ventricular hypertrophy, n | 5 |
| Mean left ventricular wall diameter, mm | 13.4 |

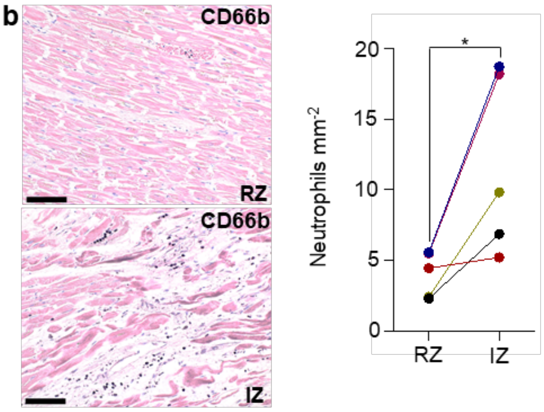


**Supplemental Figure 3: Characteristics of human AMI patients recruited for the study.** **a**, Flowchart of patient selection, individual cause of death, exclusion criteria and baseline characteristics of patients diagnosed with acute MI. **b**, Quantification of intravascular and extravascular $CD66^{+}$ neutrophils in non-infarcted remote zone (RZ) and infarcted zone (IZ) in myocardial biopsies of MI humans. n = 5. Data are mean ± s.e.m. Two-sided Mann Whitney test.

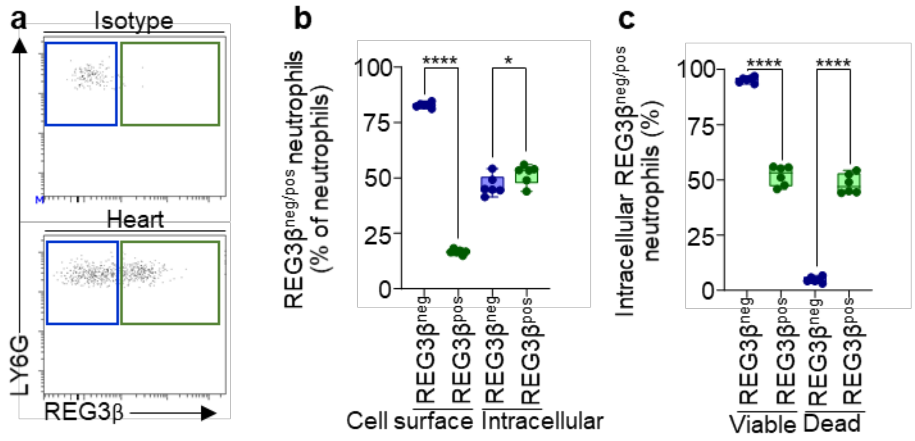


**Supplemental Figure 4: Increased death of neutrophils after MI is associated with increased uptake of REG3β. a**, Representative flow cytometric dot plots of intracellular REG3β$^{neg}$ (blue) and REG3β$^{pos}$ (green) neutrophil subsets obtained from WT hearts 2 days after MI. **b**, Flow cytometric quantification of cell surface and intracellular REG3β$^{neg}$ and REG3β$^{pos}$ neutrophils in % of neutrophils of WT hearts 2 days after MI. n = 6. **c**, Flow cytometric quantification of viable and dead cells in % of intracellular REG3β$^{neg}$ and REG3β$^{pos}$ neutrophils determined by Live/Dead Zombie dye. n = 6. Data are mean ± s.e.m. One-way ANOVA followed by Sidak's multiple comparison test. All experiments were conducted with male mice.

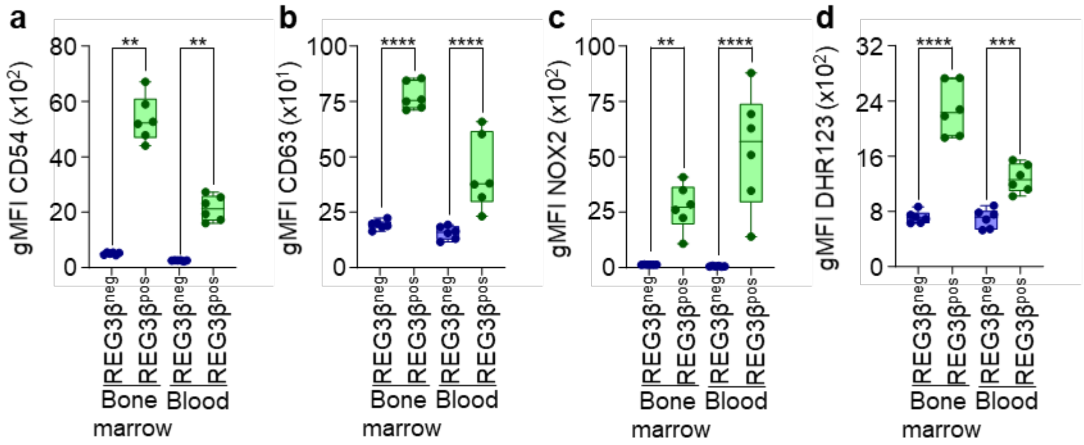


**Supplemental Figure 5: Reg3β-binding identifies hyperactivated neutrophils irrespective of the niche. a–d**, Geometric mean fluorescence intensity (gMFI) of CD54 (a), CD63 (b), NOX2 (c), and DHR123 (d) of REG3$\beta^{neg}$ (blue) and REG3$\beta^{pos}$ (green) neutrophils from bone marrow and blood of wild-type (WT) mice 2 days after MI. n = 6. Data are mean ± s.e.m. Kruskal-Wallis 1-way ANOVA followed by Dunn's multiple comparison test (**a**) and one-way ANOVA followed by Sidak's multiple comparison test (**b–d**). All experiments were conducted with male mice.

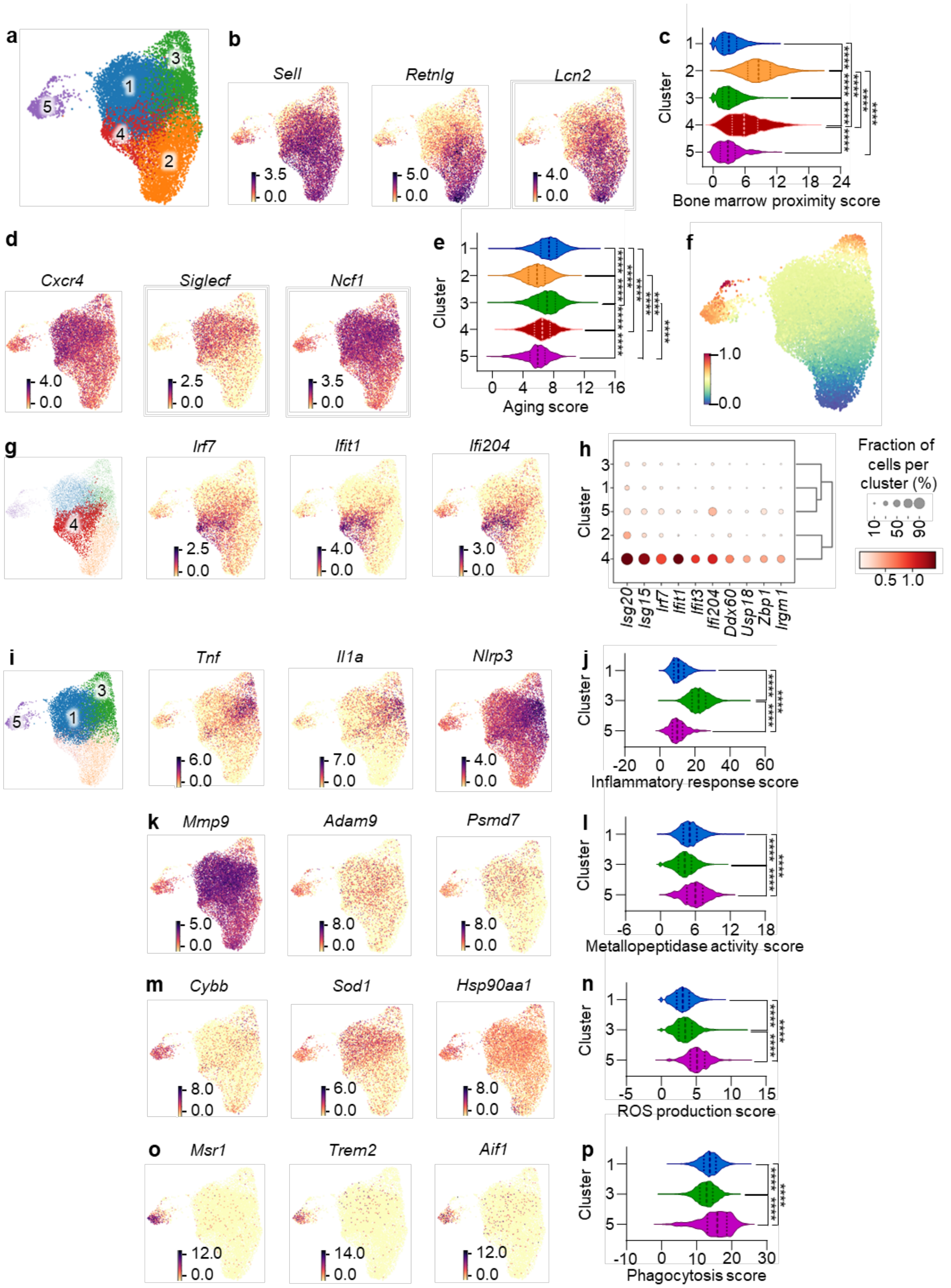


**Supplemental Figure 6: scRNA-seq analysis of REG3β$^{neg}$ and REG3β$^{pos}$ neutrophils in the heart after MI. a**, Uniform manifold approximation and projection (UMAP) of 15994 cardiac tissue neutrophils. scRNA-seq data were obtained from WT hearts at day 2 after MI. **b**, Gene expression pattern of *Sell*, *Retnlg*, and *Lcn2* projected on the UMAP plot. **c**, Violin plot of bone marrow proximity scores for each cluster. **d**, Gene expression pattern of *Cxcr4*, *Siglecf*, and

*Ncf1* projected on UMAP plots. **e**, Violin plot of aging score for each cluster. **f**, Pseudotime UMAP. **g**, Gene expression pattern of *Irf7*, *Ifit1*, and *Ifi204* projected on UMAP plots. **h**, Dot plot showing scaled expression of selected signature genes for cluster 4, colored by the average expression of each gene in each cluster scaled across all clusters. Dot size represents the percentage of cells in each cluster. **i**, Gene expression pattern of *Tnf*, *Il1a*, and *Nlrp3* projected on UMAP plots. **j**, Violin plot of inflammatory response score for cluster 1–3. **k**, Gene expression pattern of *Mmp9*, *Adam9*, and *Psmd7* projected on UMAP plots. **l**, Violin plot of metallopeptidase activity score for cluster 1–3. **m**, Gene expression pattern of *Cybb*, *Sod1*, and *Hsp90aa1* projected on UMAP plots. **n**, Violin plot of reactive oxygen species (ROS) production score for cluster 1–3. **o**, Gene expression pattern of *Msr1*, *Trem2*, and *Aif1* projected on UMAP plots. **p**, Violin plot of phagocytosis score for cluster 1–3. Data are mean ± s.e.m. Kruskal-Wallis 1-way ANOVA followed by Dunn's multiple comparison test. All experiments were conducted with male mice.

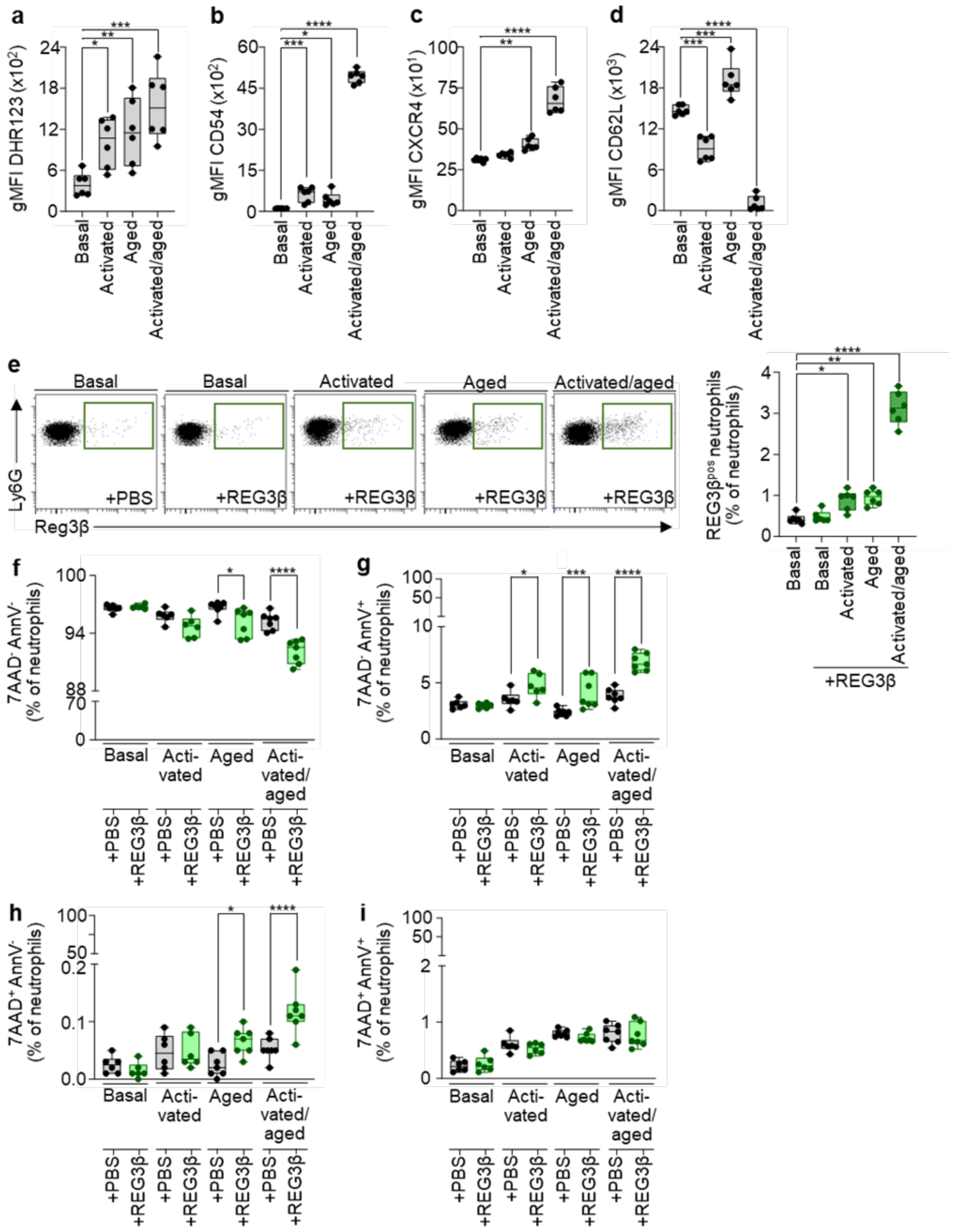


**Supplemental Figure 7: Activation of neutrophils enables binding and cytotoxicity of REG3β. a–d**, Geometric mean fluorescence intensity (gMFI) of DHR123 (a) CD54 (b), CXCR4 (c), and CD62L (d) of bone marrow neutrophils after administration of PBS (basal) and lipopolysaccharide (LPS) for 2 hours (activation): administration of PBS (aged) and LPS for 24 hours (activated/aged). n = 6 for all groups. **e**, Representative flow cytometric dot plots and quantification of REG3β$^{pos}$ (green) neutrophils under basal, activated, aged, and activated/aged conditions followed by treatment with REG3β (100ng ul$^{-1}$). PBS served as negative control. n = 6 for all groups. **f–i**, Flow cytometric quantification of viable and dead neutrophils under basal, activated, aged, and activated/aged conditions followed by treatment with REG3β (100ng ul$^{-1}$) and 7-Aminoactinomycin D (7AAD) and Annexin V (AnnV) staining. PBS was used as negative control. n = 6 for all groups. Data are mean ± s.e.m. One-way ANOVA followed by Dunnett's multiple comparison test (**a–d**) and one-way ANOVA followed by Sidak's multiple comparison test (**f–i**). All experiments were conducted with male mice.

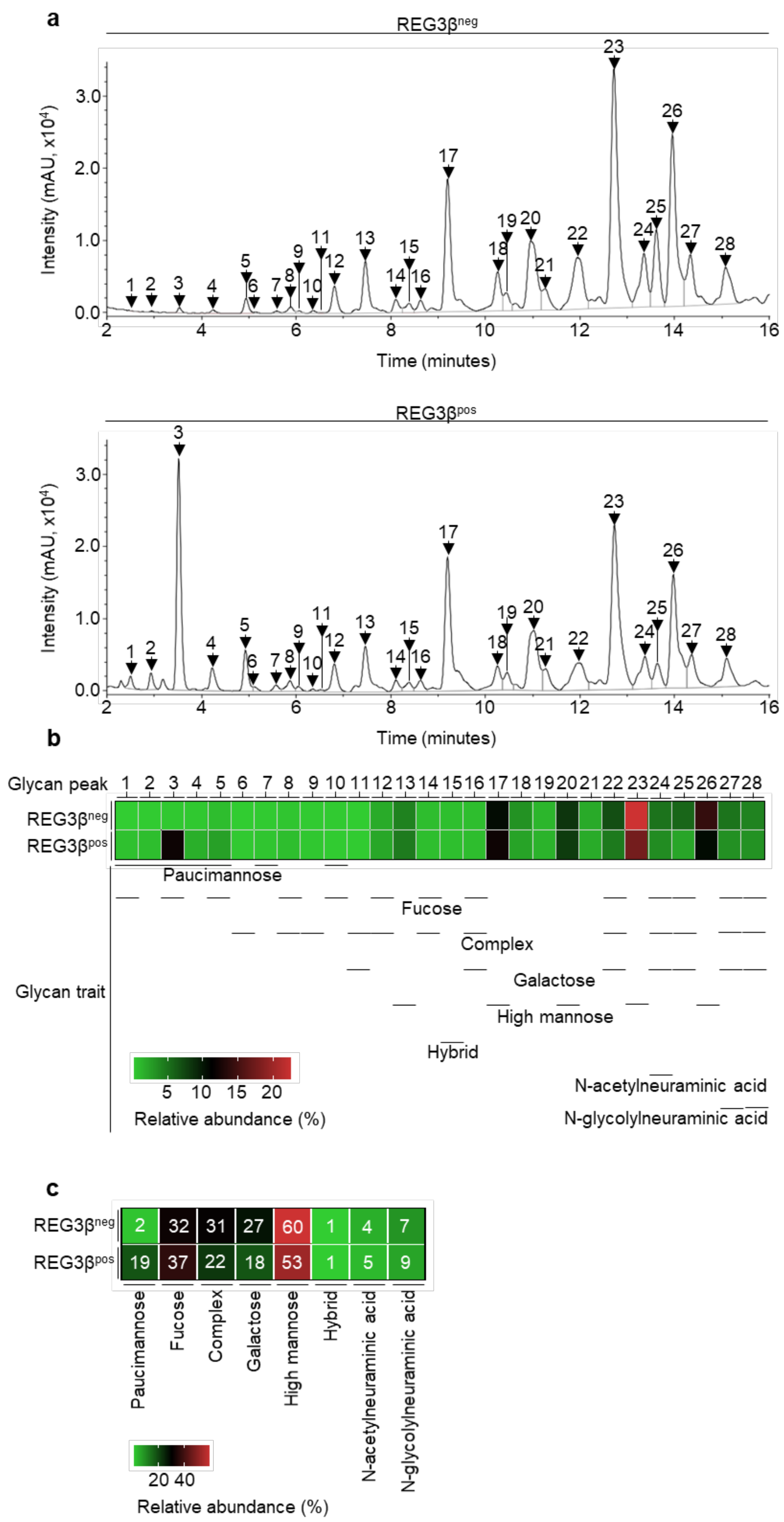
a
REG3βneg
REG3βpos
Intensity (mAU, x10⁴)
Time (minutes)
b
Glycan peak
Glycan trait
Paucimannose
Fucose
Complex
Galactose
High mannose
Hybrid
N-acetylneuraminic acid
N-glycolylneuraminic acid
Relative abundance (%)
c
REG3βneg 2 32 31 27 60 1 4 7
REG3βpos 19 37 22 18 53 1 5 9
Relative abundance (%)

**Supplemental Figure 8: Cell surface N-glycan analysis of REG3$\beta^{neg}$ and REG3$\beta^{pos}$ neutrophils.** **a**, Cell surface N-glycan analysis of sorted REG3$\beta^{neg}$ and REG3$\beta^{pos}$ peritoneal neutrophils by ultra-high-performance liquid chromatography based on hydrophilic interactions with fluorescent detection (HILIC-UHPLC-FLD) from 4 WT mice. 28 glycans were identified. **b**, Relative abundance of glycan peaks 1–28 in REG3$\beta^{neg}$ and REG3$\beta^{pos}$ peritoneal neutrophils and assignment to glycan traits based on shared structural motifs. **c**, Relative abundance of glycan traits in REG3$\beta^{neg}$ and REG3$\beta^{pos}$ peritoneal neutrophils. All experiments were conducted with male mice.

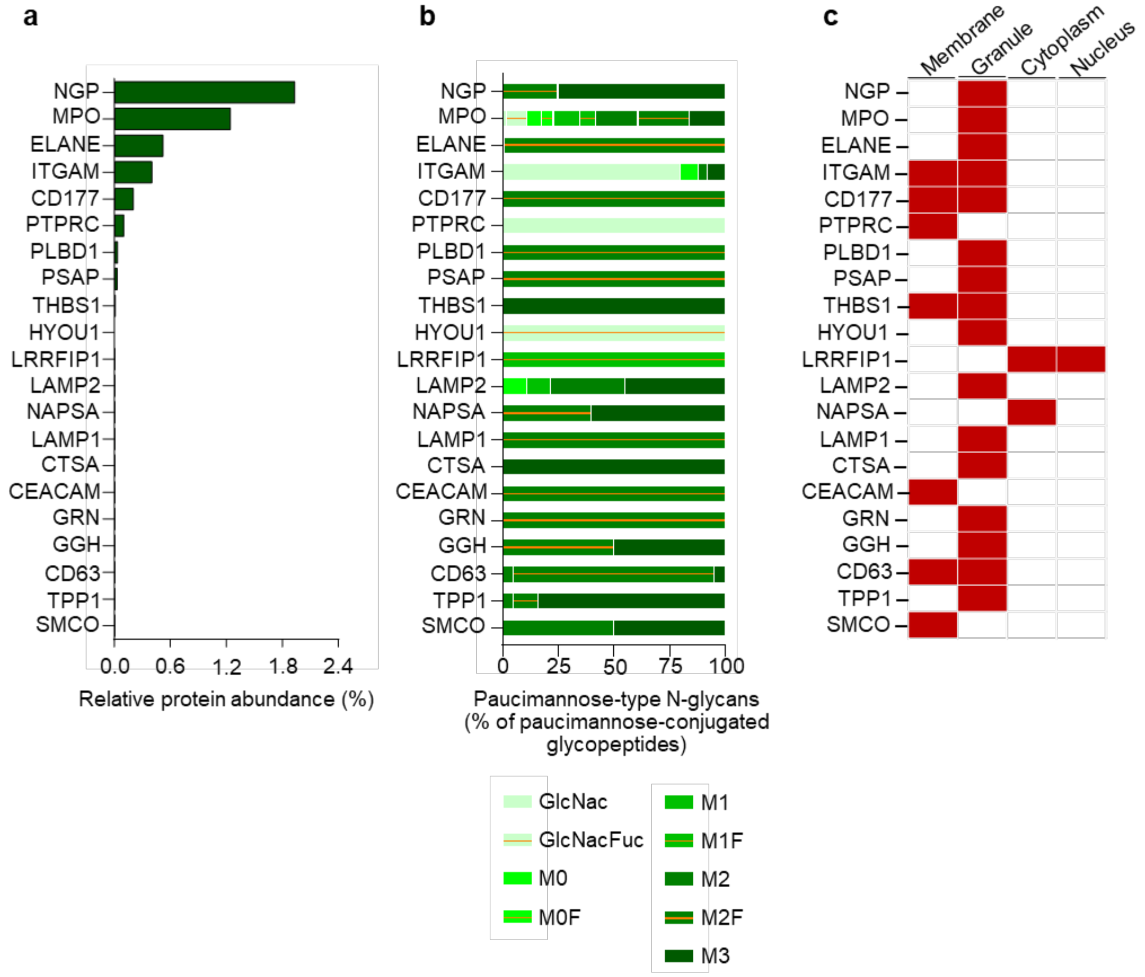


**Supplemental Figure 9: Subcellular distribution of paucimannosilated proteins and different glycoconjugates in murine neutrophils.** **a**, Relative abundance of 21 paucimannosidic proteins related to the murine neutrophil proteome detected by LC-MS/MS. **b**, Relative glycoforms of each paucimannosidic protein. **c**, Subcellular distribution of paucimannosylated proteins as described in literature and neXtprot (www.nextprot.org). Abbreviations: GlcNac - N-Acetylglucosamine, GlcNacFuc - fucosylated GlcNac, M - Mannose, F - Fucose. Data are mean ± s.e.m. Analysis was conducted with male mice.

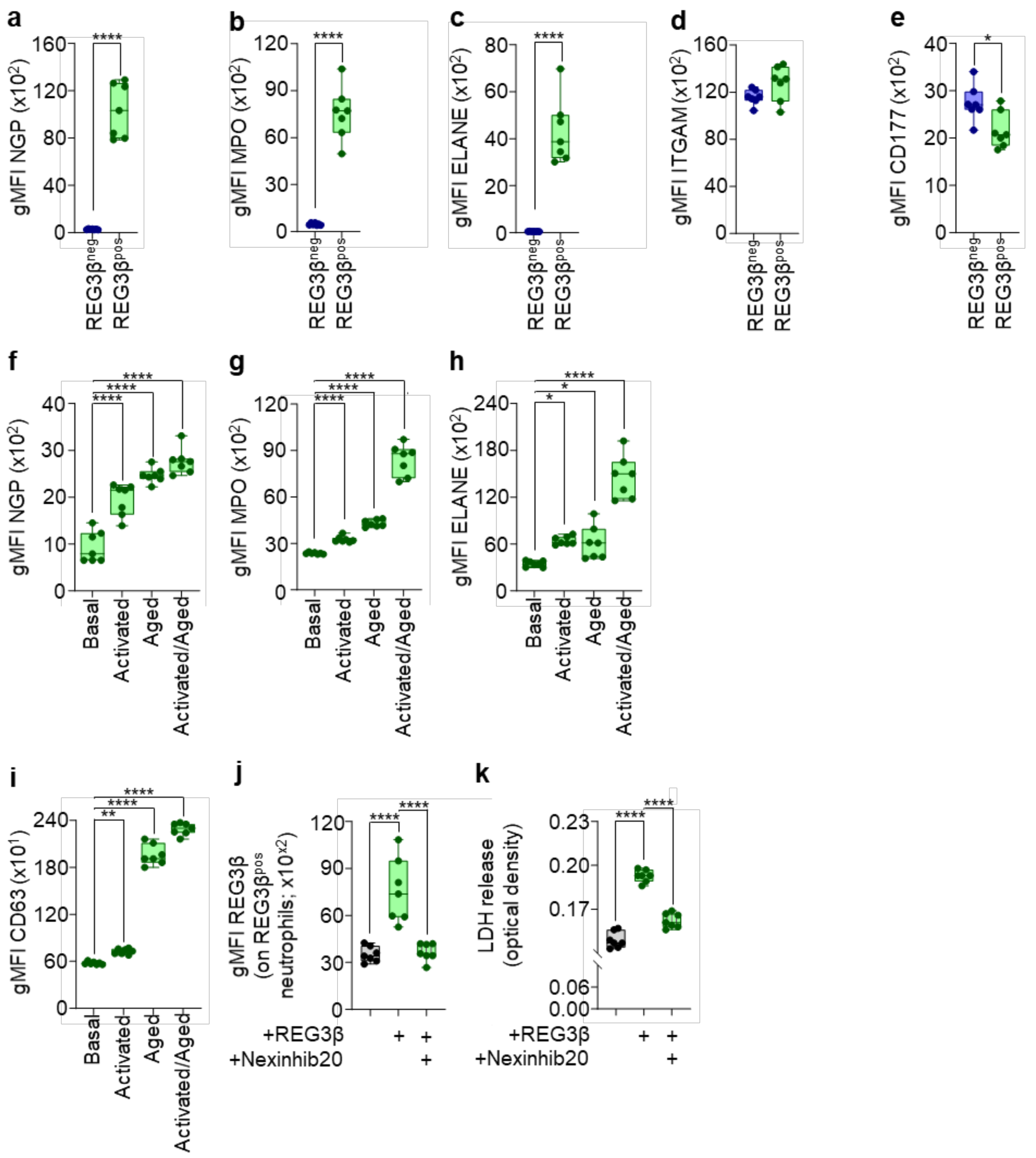


**Supplemental Figure 10: Increased presence of granule-derived, paucimannosilated proteins on the surface of REG3β$^{pos}$ neutrophils. a–e**, Geometric mean fluorescence intensity (gMFI) of neutrophilic granule protein (NGP, a), myeloperoxidase (MPO, b), neutrophil elastase (ELANE, c), integrin alpha M (ITGAM, d) and CD177 antigen (CD177, e) on REG3β$^{neg}$ and REG3β$^{pos}$) neutrophils from WT hearts 2 days after MI. n = 7. **f–j**, gMFI of NGP (f), MPO (g), and ELANE (h) of bone marrow neutrophils after administration of PBS (basal) and lipopolysaccharide (LPS) for 2 hours (activation); administration of PBS (aged) and LPS administration for 24 hours (activated/aged). n = 7 for all groups. **i**, gMFI of CD63 of basal, activated, aged, and activated/aged bone marrow neutrophils. n = 7 for all groups. **j**, gMFI of REG3β on REG3β$^{pos}$ bone marrow derived neutrophils pretreated with Nexinhib20 (10μM) for

30 minutes following by treatment with REG3β (100 ng $ml^{-1}$) for 15 minutes. DMSO served as control. n = 7. **k**, Lactate dehydrogenase (LDH) release of activated bone marrow derived neutrophils treated with Nexinhib20 (10μM) for 30 minutes following administration of REG3β (100 ng $ml^{-1}$) for 15 minutes. DMSO served as control. n = 7. Data are mean ± s.e.m. Two-sided unpaired t tests (**a–e**), one-way ANOVA followed by Dunnett's multiple comparison test (**f–i**), one-way ANOVA followed by Sidak's multiple comparison test (**j**, **k**). All experiments were conducted with male mice.

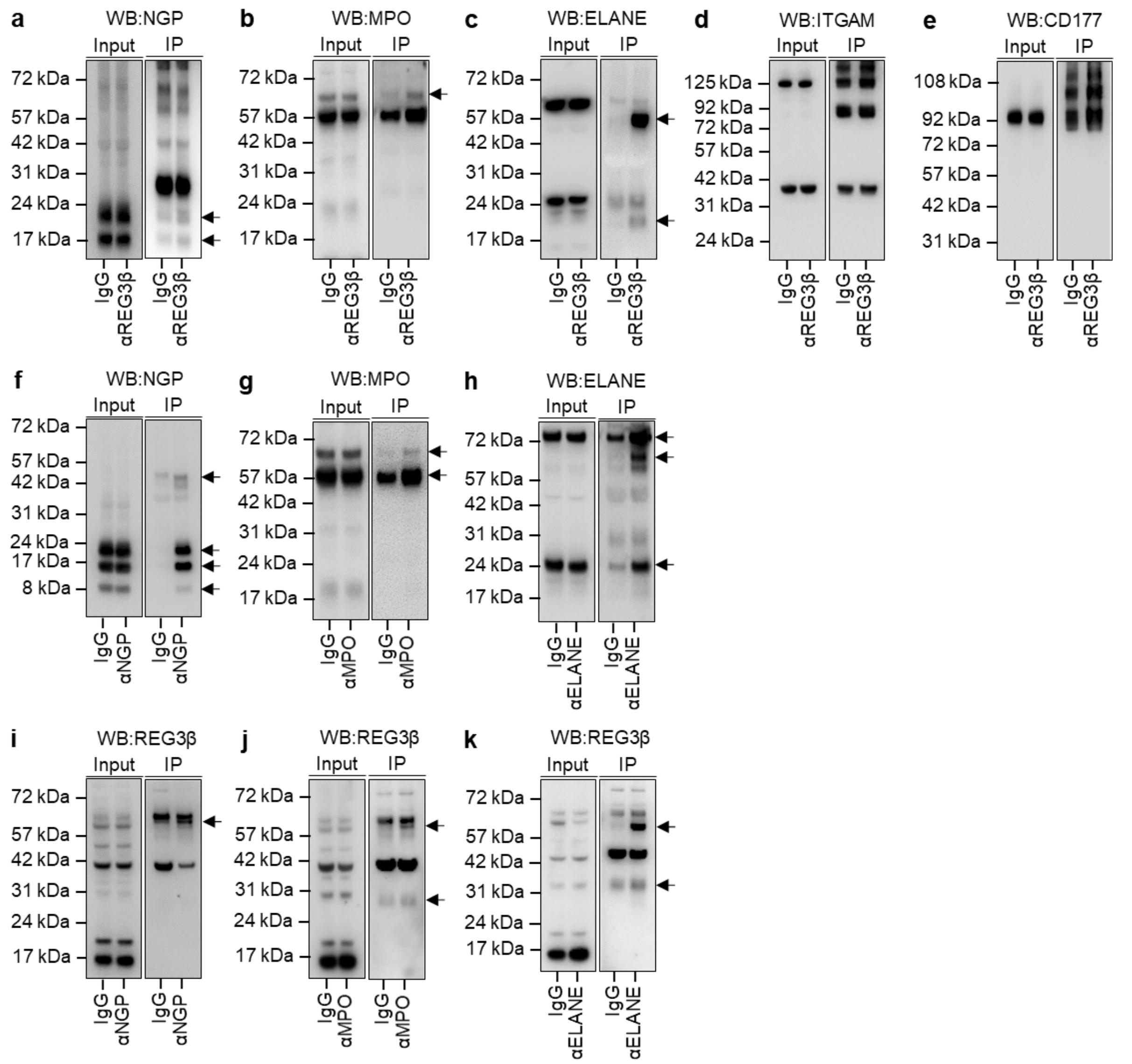


**Supplemental Figure 11: REG3β interacts with granule-derived paucimannose-conjugated proteins on the surface of neutrophils. a–k**, Immunoblot analysis of input and cell surface co-immunoprecipitated samples from peritoneal neutrophils using antibodies against REG3β, NGP, MPO, ELANE, ITGAM, and CD177. Isotype controls (IgG) were used as controls. Arrows indicate bands co-precipitated with the primary antigen.
